# Gene structure-based homology search identifies highly divergent putative effector gene family

**DOI:** 10.1101/2021.09.24.461719

**Authors:** David L. Stern, Clair Han

## Abstract

Homology of highly divergent genes often cannot be determined from sequence similarity alone. For example, we recently identified in *Hormaphis cornu* a family of rapidly evolving *bicycle* genes, which encode novel proteins implicated as plant gall effectors. Sequence similarity search methods yielded few putative *bicycle* homologs in other species. Coding sequence-independent features of genes, such as intron-exon boundaries, often evolve more slowly than coding sequences, however, and can provide complementary evidence for homology. We found that a linear logistic regression classifier using only structural features of *bicycle* genes identified many putative *bicycle* homologs in other species. Independent evidence from sequence features and intron locations supported homology assignments. To test the potential roles of *bicycle* genes in other aphids, we sequenced the genome of a second gall-forming aphid, *Tetraneura ulmi*, and found that many *bicycle* genes are strongly expressed in the salivary glands of the gall forming foundress. In addition, *bicycle* genes are strongly overexpressed in the salivary glands of a non-gall forming aphid, *Acyrthosiphon pisum*, and in the non-gall forming generations of *Hormaphis cornu*. These observations suggest that Bicycle proteins may be used by multiple aphid species to manipulate plants in diverse ways. Incorporation of gene structural features into sequence search algorithms may aid identification of deeply divergent homologs, especially of rapidly evolving genes involved in host-parasite interactions.

## Introduction

One challenge that is faced by evolutionary and functional genetic studies is that many genes have been identified as “orphan” or lineage-specific genes because homologous sequences cannot be identified outside of a limited taxonomic range. Many lineage-specific genes may not be truly novel, however, since sequence divergence can cause homologs to become undetectable by sequence-search methods (Vakirlis et al., 2020; Weisman et al., 2020). Identifying such extremely divergent homologs remains a significant bioinformatic challenge and limits functional inferences derived from homology (Loewenstein et al., 2009).

Genes involved in host-parasite systems often evolve extremely divergent sequences as a result of genetic conflict (Eizaguirre et al., 2012; Paterson et al., 2010). Aphids and their host plants represent one such antagonistic pair. Aphids are small insects that feed by inserting their thin mouthparts (stylets) into the phloem vessels of plants to extract nutrients (Dixon, 1978). Like many herbivorous insects, aphids introduce effector molecules into plant tissues to manipulate the physiology and development of plants to the insects’ advantage (Elzinga and Jander, 2013; Hogenhout and Bos, 2011; Mutti et al., 2008; Rodriguez et al., 2017). For example, aphids introduce calcium binding proteins that prevent the plant’s ability to block phloem cell transport in response to phloem vessel damage (Will et al., 2007). It is believed that aphids introduce a wide range of effector proteins and that these molecules contribute to the debilitating effects of multiple aphid species on plants, which imposes significant financial damage on most major agricultural crops (Schaible et al., 2008).

In an extreme form of aphid manipulation of plant physiology and development, some aphid species induce novel plant organs, called galls, which provide the aphids with a ready food source and with protection from the elements and from natural enemies (Giron et al., 2016; Mani, 1964; Shorthouse et al., 2005). Galls can be significant nutrient sinks (Burstein et al., 1994), demonstrating that galling aphids induce both local and long-range changes to plant physiology.

We reported recently that the genome of one species of galling aphid encodes 476 related genes encoding diverse, novel, presumptive effector proteins and many of these proteins contain two cysteine-tyrosine-cysteine motifs (CYC) (Korgaonkar et al., 2021). These genes were therefore named *bicycle* (bi-CYC-like) genes. A genome-wide association study revealed that variation in one *bicycle* gene, called *determinant of gall color* (*dgc*), was strongly associated with a red versus green gall color polymorphism and this genetic polymorphism was associated with strong downregulation of *dgc* in aphids and strong upregulation of a small number of genes involved in anthocyanin production in plant galls. In addition, the majority of *bicycle* genes are strongly upregulated in the salivary glands specifically of the aphid generation that induces galls (the fundatrix). Together with genetic evidence that *dgc* regulates the gall phenotype, the enrichment of *bicycle* genes expressed in fundatrix salivary glands suggests that many *bicycle* genes contribute to gall development or physiology.

The primary amino-acid sequences of Bicycle proteins have evolved rapidly, apparently in response to extremely strong positive selection (Korgaonkar et al., 2021). In preliminary studies, we attempted to identify *bicycle* homologs in other insect species using sequence similarity algorithms such as *BLAST* (Altschul et al., 1990) and *hmmer* (Eddy, 2011; Johnson et al., 2010), but identified few putative homologs. It was therefore unclear if *bicycle* genes are a recently evolved family of genes or an ancient family that has evolved highly divergent coding sequences.

Extremely divergent homologs have been identified previously using features of gene structures that evolve more slowly than the coding sequence, such as exon sizes and intron positions (Bazan, 1991; Betts, 2001; Brown et al., 1995), and these characteristics of genes have also proven to be valuable for phylogeny reconstruction (Rokas et al., 1999; Rokas and Holland, 2000; Telford and Copley, 2011; Venkatesh et al., 1999). We noted previously that *bicycle* genes have unusual gene structures containing many micro-exons (Korgaonkar et al., 2021) and we considered the possibility that *bicycle* gene structure may be more conserved than *bicycle* gene sequences. To explore the evolutionary history of *bicycle* genes, we therefore sought a method that would allow identification of highly divergent *bicycle* gene homologs that did not rely on sequence similarity. To accomplish this, we built a logistic regression classifier based only on structural features of *bicycle* genes. We found that this classifier was very accurate, despite the fact that it does not include any sequence information. This classifier identified many highly-divergent *bicycle* homologs in *H. cornu* that could not be identified by sequence similarity search methods, including genes that encode proteins that do not include the previously canonical CYC motif. In addition, the classifier identified many highly divergent candidate *bicycle* homologs in all aphids, phylloxerans, and scale insects we studied. We did not detect any putative homologs in three progressively basal outgroups. Multiple sequence alignment of these putative homologs revealed that many contain N-terminal signal sequences and CYC motifs, consistent with their assignment as *bicycle* homologs. In addition, gene-structure aware sequence alignment revealed multiple apparently shared intron boundaries between putative homologs that share little obvious sequence similarity. In addition, putative *bicycle* homologs are highly enriched in the salivary gland mRNA of two gall forming and one non-gall forming aphid. All of this evidence supports the hypothesis that *bicycle* gene homologs encode effector proteins in gall-forming and non-gall-forming aphids and also in phylloxerans and scale insects.

## Results

### Sequence-based homology searches find few bicycle genes outside of H. cornu

*Bicycle* genes in *H. cornu* were identified originally as a subset of previously unannotated genes that were enriched in the salivary gland mRNA of the gall-inducing foundress (Korgaonkar et al., 2021). *Bicycle* genes were found to be the single largest group of such genes that shared sequence similarity. These 476 genes are extremely divergent from one another at the amino-acid sequence level and appear to have evolved rapidly due to positive natural selection. Using sequence-based homology search methods *BLAST* and *hmmer*, we identified a few additional candidate *bicycle* paralogs in *H. cornu* (using *BLAST*) and putative homologs in other aphid species (mainly using *hmmer*). We did not detect any candidate *bicycle* orthologs in genomes from insects of the families Phylloxeridae, Coccoidea, Psylloidea, Aleyrodoidea, and Fulgoroidea (Figure 1).

**Figure 1.**
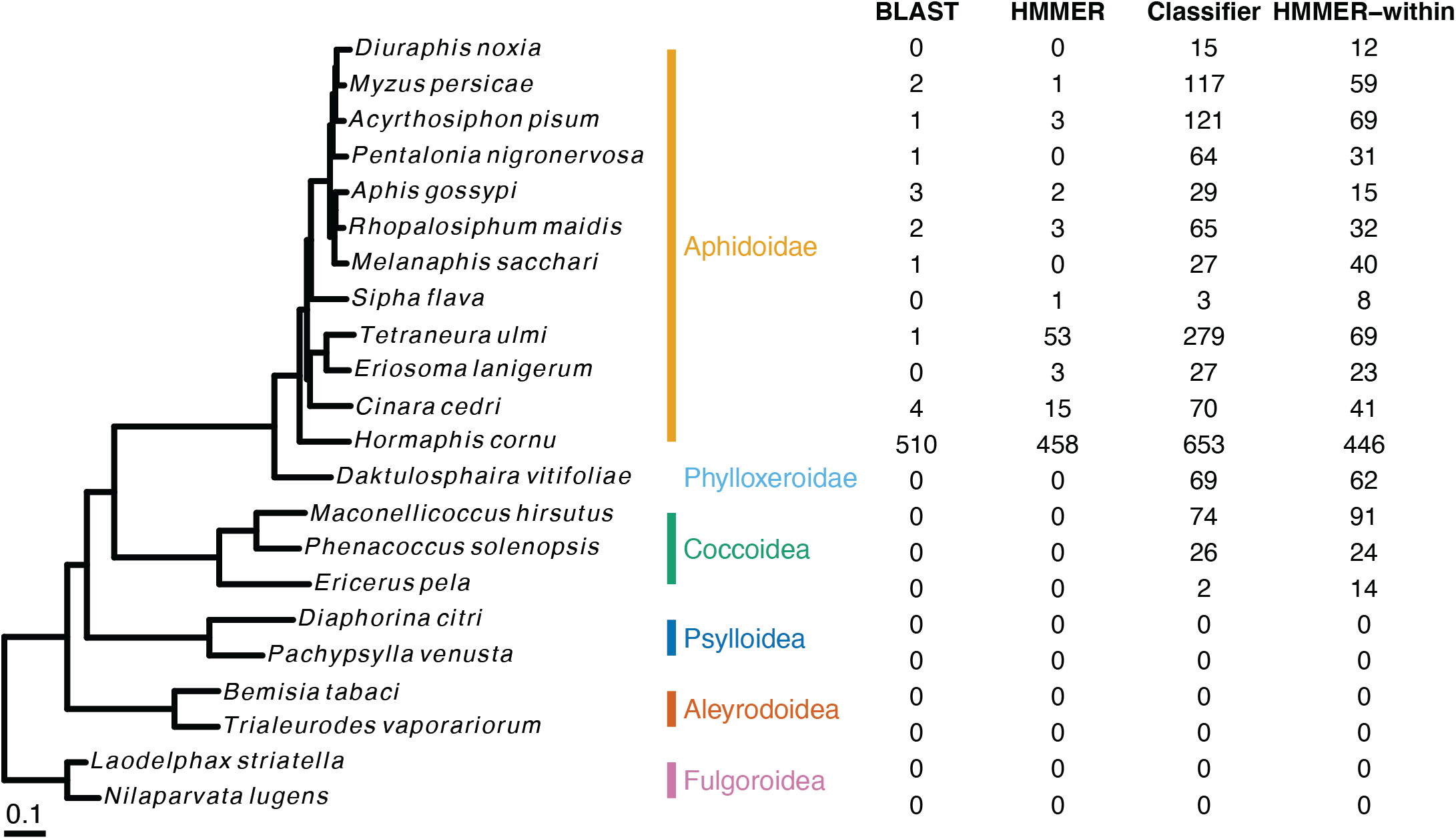
The gene-structure based classifier detects *bicycle* genes in genomes of aphids, phylloxerids, and coccids. A whole-proteome phylogeny of the species studied is shown on the left. Family membership is shown in color beside the species names. The number of candidate *bicycle* homologs detected by *BLAST*, *hmmer*, and the gene-structure based classifier is shown to the right of the phylogeny. The number of candidate *bicycle* homologs detected by profile *hmmer* starting with the *bicycle* homologs identified by the gene-structure based classifier within each species is shown in the far right column.

### A classifier based on only gene structural features can identify *bicycle* genes with high probability

Sequence-based search methods suggested that *bicycle* genes may be found in other species, but the low number of genes identified, and the incongruence between results from *BLAST* and *hmmer* suggested that sequence-based search may provide low sensitivity to detect *bicycle* homologs (Vakirlis et al., 2020; Weisman et al., 2020). We noticed, however, that while sequence divergence was high at the amino-acid level among *H. cornu bicycle* genes, non-sequence specific features of *bicycle* genes appeared to be relatively well conserved. For example, *bicycle* genes contain a large number of unusually small internal exons (Korgaonkar et al., 2021), making them clear outliers to most genes in the *H. cornu* genome. We also observed that almost all internal exons of *bicycle* genes start with codon position 2, that is, they exhibit exon phase 2 (Figure S1A), which is an extremely different distribution of exon phases compared with other genes in the genome (Figure S1B). Genes containing many exons of similar size all with the same phase are rare in most genomes (Ruvinsky and Watson, 2007).

We hypothesized that some aspects of *bicycle* gene structure may evolve more slowly than the primary protein sequences and thus provide an evolutionary signal to detect distantly related *bicycle* homologs (Aguiar et al., 2015). For example, the pattern of exon phases can allow identification of highly-divergent homologous genes (Roy and Gilbert, 2005; Ruvinsky and Watson, 2007). Development of a full model to characterize gene structure evolution is beyond the scope of this paper, but we explored whether a linear logistic regression classifier could identify *bicycle* homologs based on the following structural features of each gene: length of gene in genome, from start to stop codons; first and last exon length; internal exon mean length; length of the sum of codons; and number of internal exons that start in phase 0, 1, or 2. Since almost all internal *bicycle* exons are of phase 2, there is no additional information from the order of exon phases in *bicycle* genes, and we therefore simply counted the number of exons of each phase.

Characterized *bicycle* genes have an average of approximately 17 and a minimum of 5 exons (Figure S1C), which is strongly different from the genome-wide background (Figure S1D), and Bicycle protein lengths are well explained by total exon number (c.f. Figures S1E and S1F). We excluded all genes with zero or one internal exon to accurately summarize internal exon mean lengths and number of internal exons that start in each of the three phases. Thus, if any *bicycle* genes have evolved to have three or fewer exons, then they could not be detected with this classifier.

The classifier exhibited extremely good performance with all data partitions (Figure S2A) and, therefore, a full model was constructed using all *bicycle* and annotated genes. (Non-annotated genes were excluded from model training and validation because we hypothesized that this set may include additional *bicycle* homologs.) The classifier categorized the vast majority of genes as either non-*bicycle* or *bicycle* genes with high probability (Figure S2B). To decide on a probability cutoff to identify putative *bicycle* homologs, we considered the trade-off between precision (the proportion of true positives among both true and false positives) and recall (the proportion of true positives among true positives and false negatives). We chose a model probability cutoff that equalized precision and recall (P/R = 1), which yielded precision and recall values of 0.99 (Figure S2C, D).

To determine whether all gene features were required for accurate classification, we removed one factor at a time, retrained the model, and calculated precision and recall values using the same threshold as the full model. We found that no single factor was critical for model performance (Figure S2E), although removal of the number of mode 2 exons resulted in the strongest decrement in model performance, resulting in precision and recall values of 0.98 and 0.96, respectively.

To determine whether any single predictor was sufficient to identify *bicycle* genes with high confidence, we calculated precision and recall values for linear models using one predictor at a time (Figure S2F). Mean internal exon length alone performed best (Figure S2F), consistent with the observation that *bicycle* genes contain an unusually large number of small exons (Figure S3). The number of internal exons in mode 2 also displayed some predictive power on its own (Figure S2F). No other predictors alone had substantial predictive power for discriminating *bicycle* genes from the background (Figure S2F). We observed some level of correlation between predictor variables (Figures S2G-I), but no variables were scalar multiples of each other and we therefore employed the full model using all predictors to search for new *bicycle* genes.

### Additional candidate *bicycle* genes can be found in many other species

To search for candidate *bicycle* genes in other species, we applied the classifier to genes from the genomes of twenty-two species spanning the Aphidoidea and species from the five most closely related families, the Phylloxeroidea, Coccoidea, Psylloidea, Aleyrodoidea, and Fulgoroidea (Figure 1). Multiple *bicycle* genes were identified in all Aphidoidea, Phylloxeroidea, and Coccoidea we studied and the *bicycle* genes identified by the classifier encompass 98.2-100% of the *BLAST* identified bicycle genes and 86.8-100% of the *HMMER* identified *bicycle* genes in all species with more than 3 *bicycle* genes. This concordance of methods supports the inference that (1) the classifier has identified *bicycle* homologs and (2) the classifier has higher power than sequence-based approaches to identify *bicycle* genes. As expected from the features of the classifier, the candidate *bicycle* genes in all species exhibited the unusual feature of being encoded by genes containing many micro- exons (Figure S3).

While the classifier exhibits substantial power to detect highly divergent *bicycle* genes across species, the classifier might fail to identify *bicycle* genes with gene structures that are divergent from the *bicycle* genes originally identified in *H. cornu* that were used to train the classifier. In addition, the genomes of many species studied here are highly fragmented, and some true *bicycle* genes may have been annotated with a subset of their exons. We therefore performed profile *hmmer* search within each species starting with the classifier-identified *bicycle* genes in each species (last column in Figure 1). The *hmmer* search identified additional candidate *bicycle* genes in some species, suggesting that the classifier underestimates the number of *bicycle* genes in some species.

### Candidate bicycle genes in H. cornu are highly expressed and share structural homology

When we applied the classifier to all genes in the *H. cornu* genome, we identified four false negative and 181 “false positive” genes. Since we chose a probability cutoff that should report an approximately equal number of false negatives and false positives, we hypothesized that the vast majority of “false positives” are newly discovered *bicycle* homologs. We clustered these genes together with the original *bicycle* genes based on sequence similarity (Figure 2A), and found that 528 represented a single group of related genes (Cluster 2, Figure 2A) that all share sequence similarity with, and included, the originally defined 476 *bicycle* genes (Figure 2C).

**Figure 2.**
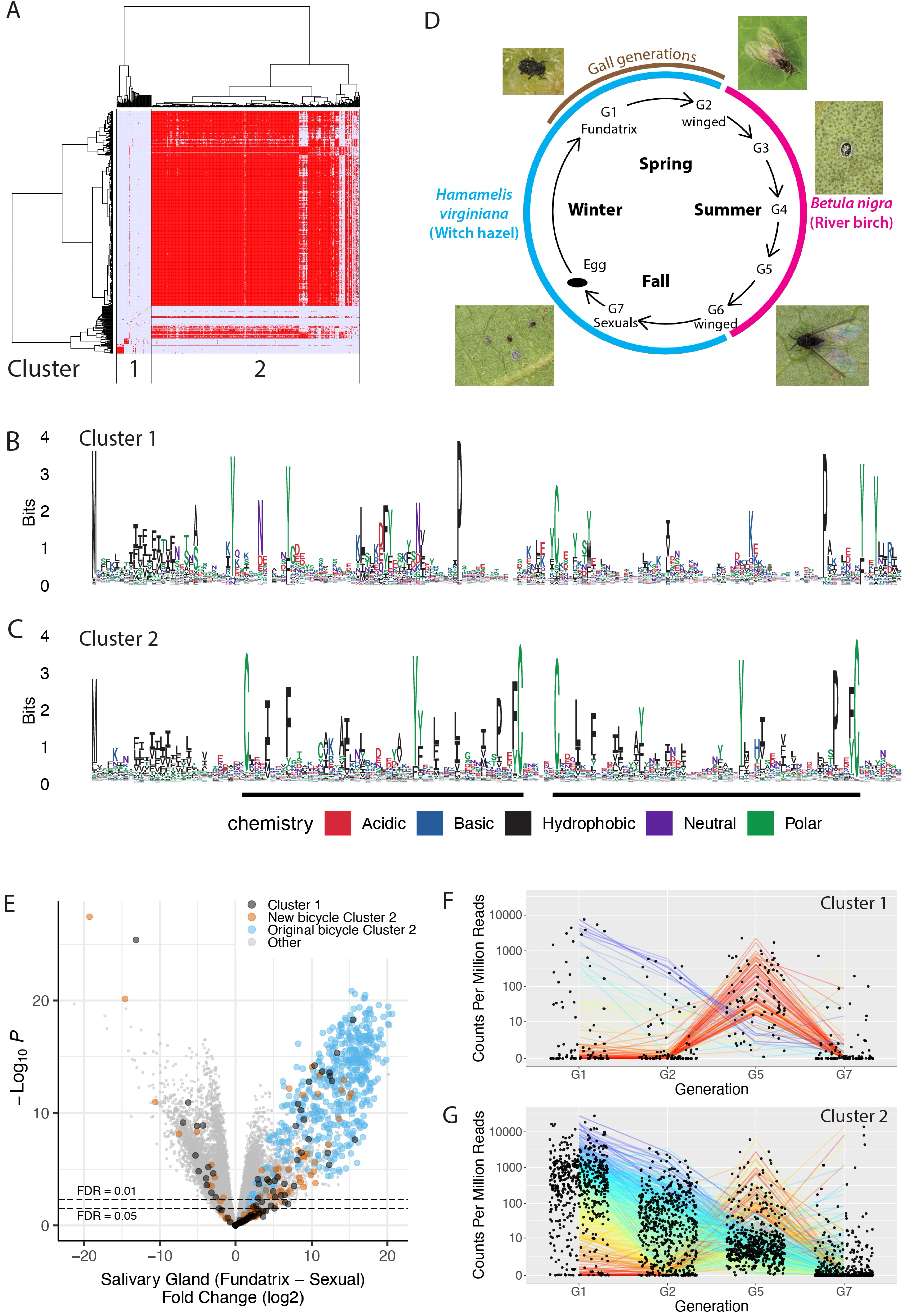
The gene-structure based classifier identifies additional CYC and non-CYC motif containing genes in the *H. cornu* genome that may contribute to the effector protein repertoire of multiple stages of the *H. cornu* life cycle. (A) Hierarchical clustering of candidate *H. cornu bicycle* homologs detected by the gene- structure based on amino-acid sequence similarity measure reveals two clusters of genes with similar sequences. Some of the genes in Cluster 1 appear to have some sequence similarity to Cluster 2 genes. (B) Genes in Cluster 1 encode proteins with N-terminal signal sequences and weak similarity to *bicycle* genes. (C) Genes in Cluster 2 encode proteins with sequence similarity to the previously described *bicycle* genes. (D) Diagram of life cycle of *H. cornu*. Individuals from different generations exhibit phenotypes that are specialized for each stage of the complex life cycle. The fundatrix (G1) is the only generation that induces a gall. (E) A volcano plot of the strength of evidence for differential expression (-log10(P value)) versus the fold-change differential expression of salivary glands isolated from fundatrices versus sexuals reveals that some of the Cluster1 (black) and 2 (orange) genes identified by the gene structure based classifier are as strongly enriched in fundatrix salivary glands as the originally described *bicycle* genes (blue). In addition, several Cluster 1 and 2 genes are more strongly enriched in salivary glands of sexuals. (F-G) Plots of gene expression levels for Cluster 1 (F) and 2 (G) genes across four generations of the life cycle reveals that while most *bicycle* homologs are most strongly expressed in fundatrix salivary glands (e.g. purple lines), some are weakly expressed in fundatrix salivary glands and then strongly expressed in salivary glands of individuals from other stages of the life cycle (e.g. red lines).

We also identified a second cluster of 129 genes that encode proteins that do not share strong sequence similarity with the originally defined Bicycle proteins and do not exhibit strong evidence for the CYC motif (Cluster 1, Figure 2A, B). Approximately 20 of these genes were strongly enriched in the salivary glands of gall-inducing foundresses, but many of these Cluster 1 genes were strongly enriched in salivary glands of other life stages, especially generations that feed on River Birch (Figure 2D- F). In addition, approximately 20 *bicycle* genes containing the CYC motif (Cluster 2) were weakly expressed in fundatrix salivary gland, but strongly expressed at other life stages (Figure 2E-G).

Like the original 476 *bicycle* genes, the new putative *bicycle* homologs likely experienced increased levels of positive selection. We found that the new putative *bicycle* homologs displayed a similar pattern to the original *bicycle* genes of elevated ratios of non-synonymous (*d_N_*) to synonymous (*d_S_*) substitutions between *H. cornu* and its closely related sister species *H. hamamelidis* compared to the genomic background (Figure S4A-C). In addition, we found that signals of selective sweeps, based on *H. cornu* population site frequencies, are enriched near both the original *bicycle* genes and the newly discovered putative *bicycle* homologs (Figure S4D).

Since some of these new putative *bicycle* homologs shared little apparent sequence similarity with the originally defined *bicycle* genes, we sought additional evidence for homology. We hypothesized that other details of gene structure not employed in the classifier, such as intron positions, might provide additional evidence of shared ancestry. We therefore performed gene-structure-aware multiple sequence alignment of the original *bicycle* genes and the new putative *bicycle* genes identified by the classifier (Gotoh, 2021).

Initial attempts to align all 653 candidate proteins were uninformative and we hypothesized that this was because these proteins display a large range of lengths and exon numbers (Figure S1C). We therefore divided Bicycle proteins of Cluster 2 into four groups based on protein length (blue dotted lines in Figure 3A). Logo plots of these four groups revealed a striking pattern, proteins contain one, two, three, or four CYC motifs (Figure 3B-E), and we therefore call these *unicycle, bicycle, tricycle,* and *tetracycle* genes, respectively. This evolutionary comparison suggests that the CYC motif is a functional unit, which can be multimerized within individual proteins.

**Figure 3.**
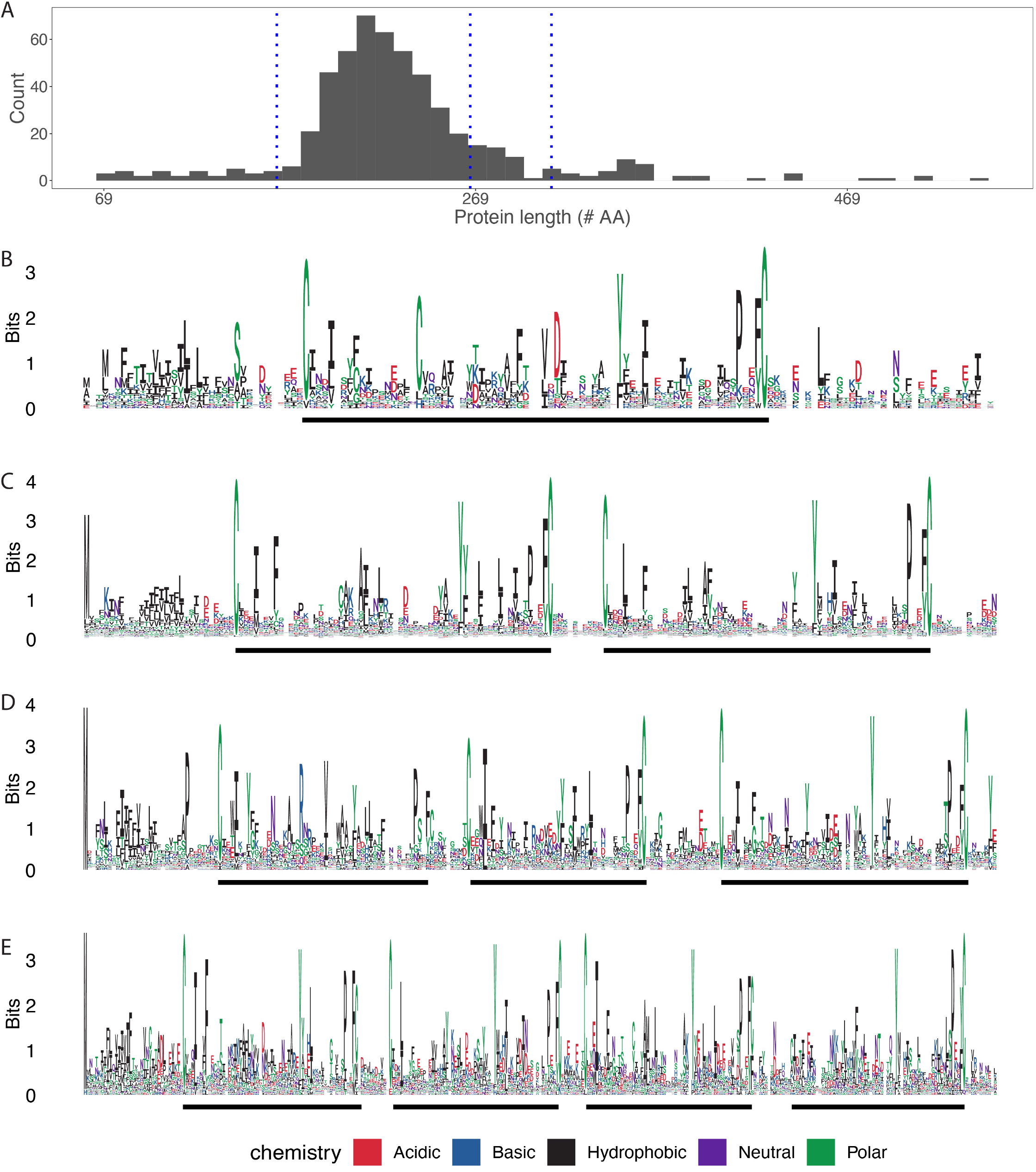
*H. cornu bicycle* genes belong to four major categories, *unicycle, bicycle, tricycle,* and *tetracycle* genes (A) The originally identified *bicycle* genes encode proteins that exhibit a wide size range, with representatives of approximately four size classes, indicated by the vertical dotted blue lines. (B-E) Logo plots of the proteins from each of the four size classes of Bicycle proteins identifies Unicycle (B), Bicycle (C), Tricycle (D), and Tetracycle (E) proteins. The CYC motifs in each protein are identified by thick black lines below each Logo plot.

We then divided *H. cornu bicycle* homologs into four length categories and performed gene-structure aware protein sequence alignments to search for shared intron position between the originally defined *bicycle* genes and the newly discovered *bicycle* putative homologs of Cluster1. We identified many evolutionarily shared intron positions between the two classes of putative homologs (Figure 5 Supplement 1A-D).

To determine whether more introns are shared between the original *bicycle* genes and newly identified putative *bicycle* genes than expected by chance, for example given an alignment of two groups of unrelated genes, we developed a metric of intron concordance and estimated the null distribution by resampling intron concordance between *bicycle* genes and three unrelated gene families (Figure S6). Intron concordance was measured as the correlation coefficient between the number of introns at each alignment position for one gene family versus the *bicycle* gene family (Figure S6A-D). Unrelated gene families were found to have intron positions distributed approximately randomly relative to *bicycle* genes with mean R close to 0 (Figure S6C, E). In contrast, newly discovered *bicycle* genes, both from *H. cornu* and from other species, displayed positive R much greater than 0 and significantly different from distributions of R values found for pairs of unrelated gene families (Figure S6E). This analysis provides statistically significant support for the conclusion that newly discovered putative *bicycle* genes share introns in the same locations more often than expected by chance between unrelated gene families, and thus, that they are likely *bicycle* gene homologs.

These results provide evidence for several conclusions. First, measures of gene structure alone can reliably identify many, but perhaps not all, divergent *bicycle* genes. Second, some *bicycle* homologs have evolved highly divergent sequences, raising the possibility that highly divergent homologs may be present in other species but are undetectable with sequence-similarity search algorithms. Third, some *bicycle* homologs are highly expressed in salivary glands of non-gall forming life stages, suggesting that some of these proteins may contribute to the effector-protein repertoire outside of the context of gall development. Given these observations, we next tested whether the classifier could identify *bicycle* homologs in a divergent galling aphid species.

### Many candidate gall effector genes in Tetraneura ulmi are highly divergent bicycle genes

There are two major clades of gall forming aphids, the Hormaphididae, to which *H. cornu* belongs, and the Pemphigidae. These two aphid families are thought to have shared a common ancestor that induced galls. Therefore, to determine whether *bicycle* genes were present in the common ancestor of gall forming aphids, we assembled and annotated the genome of *Tetraneura ulmi* (Figure S7A), a gall forming aphid belonging to the Pemphigidae (Álvarez et al., 2013). Details of the genome assembly can be found in Supplementary Methods. We annotated the *T. ulmi* genome using mRNA sequencing reads generated from salivary glands and carcasses of the fundatrix (G1) and G2 generations (Figure S7B).

We applied the *H. cornu bicycle* gene classifier to predicted genes of the *T. ulmi* genome and identified 279 candidate *bicycle* homologs (Figure 1). In contrast, sequence-based search using BLAST and *hmmer* identified 1 and 53 candidate homologs at E-value < 0.01, respectively. We clustered the 279 *T. ulmi* genes discovered by the gene-structure based classifier based on their predicted amino acid sequences (Figure 4A) and identified three clusters that included proteins with clear evidence for CYC motifs (Figure 4B-D). One cluster of 94 genes does not exhibit clear CYC motifs (Figure 4E). The extreme sequence divergence of these genes within *T. ulmi* is supported by the fact that profile *hmmer* search recovered only 69 of these original 279 genes (Figure 1). To test whether these genes likely shared a common ancestor, we performed gene-structure aware alignments of proteins with and without CYC motifs and identified more shared introns between putative homologs with and without CYC motifs (Figure S7C) than expected by chance (Figure S6E). Thus, the *T. ulmi* genome contains apparent *bicycle* genes that have become essentially unrecognizable as *bicycle* homologs at the sequence level.

**Figure 4.**
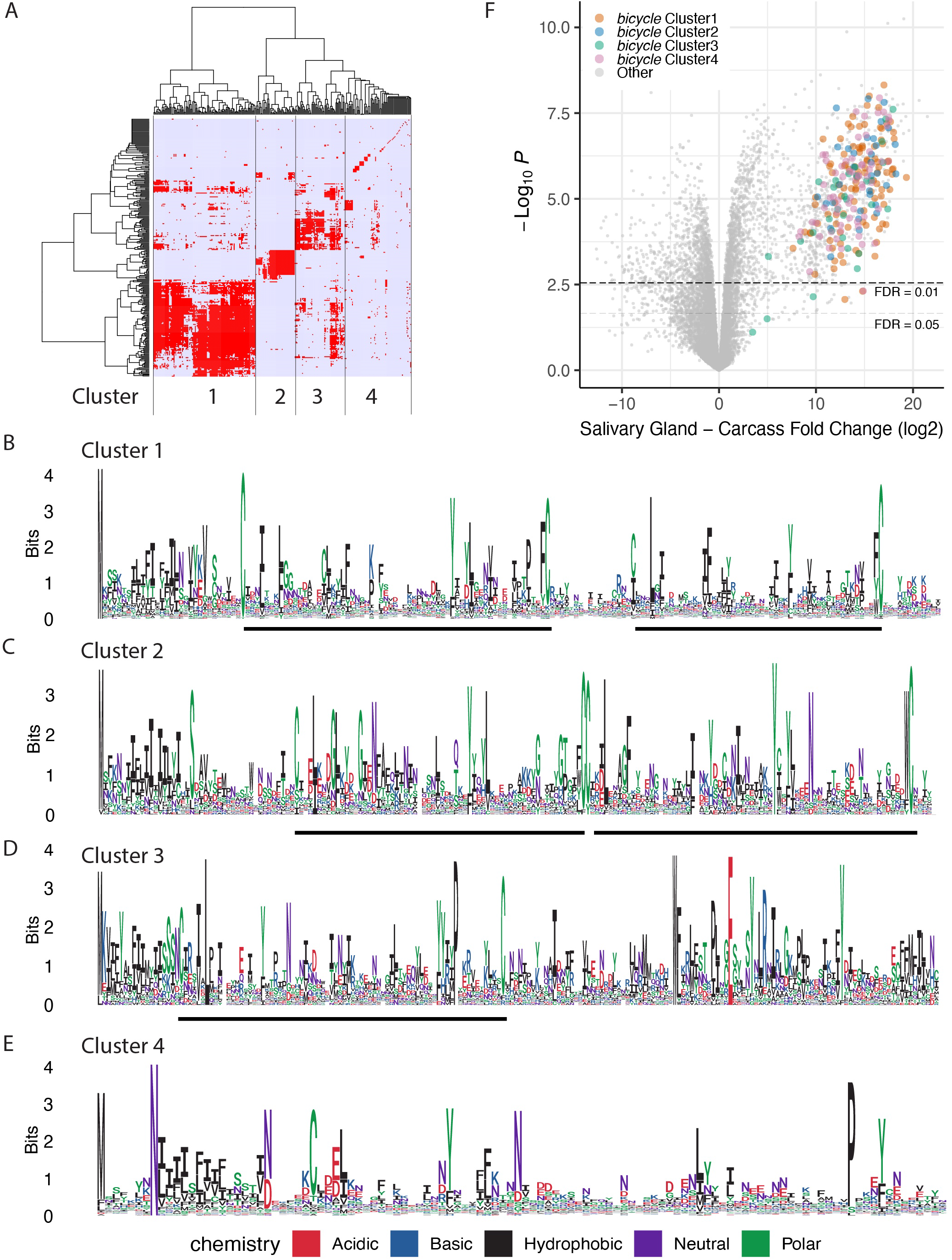
The *T. ulmi bicycle* homologs detected by the gene-structure based classifier display CYC motifs and highly divergent protein sequences. (A) Hierarchical clustering of candidate *T. ulmi bicycle* homologs based on amino-acid sequence similarity measure reveals four clusters of genes. (B-E) The four clusters of *T. ulmi bicycle* homologs display two (B, C) or one (D) CYC motif, or no clear CYC motifs (E). The cluster identities are indicated in each subplot. (A) Volcano plot of genes differentially expressed in fundatrix salivary glands versus carcass with *bicycle* genes labelled. Most of the *T. ulmi* candidate *bicycle* homologs are strongly over-expressed in the fundatrix salivary glands.

To explore whether the *T. ulmi bicycle* genes, and especially the divergent putative *bicycle* homologs, might act as effector proteins, we dissected salivary glands from fundatrices and compared mRNA expression levels in salivary glands versus carcasses. Almost all of the *bicycle* homologs detected by the classifier are amongst the most strongly over-expressed genes in the fundatrix salivary glands (Figure 4F), suggesting that they may contribute to the effector protein repertoire in *T. ulmi*.

The discovery of *bicycle* genes in *T. ulmi* using the gene-structure based classifier provides evidence that the classifier can identify putative *bicycle* homologs even when the genes cannot be identified using sequence-based homology methods and that *bicycle* genes were likely present in the common ancestor of gall-forming aphids. We therefore next tested whether the classifier can detect putative *bicycle* homologs in an aphid species that does not induce galls.

### The pea aphid (Acyrthosiphon pisum) genome includes many bicycle homologs that are candidate effector proteins

The gene-structure based classifier detected 121 putative *bicycle* homologs in the genome of *Acyrthosiphon pisum*, a species that does not induce galls (Li et al., 2019; Richards et al., 2010). In contrast, sequence-based search using *BLAST* and *hmmer* identified 1 and 3 candidate hits at E-value < 0.01, respectively. Multiple sequence alignment of the 121 *A. pisum* putative *bicycle* homologs revealed that most of these genes encode proteins with N-terminal secretion signal sequences and a single CYC motif (Figure 5A). Gene structure aware alignment identified more shared introns between *H. cornu bicycle* genes and *A. pisum* putative *bicycle* homologs (Figure S8) than expected by chance (Figure S6E), providing independent confirmation that these genes shared a common ancestor.

**Figure 5.**
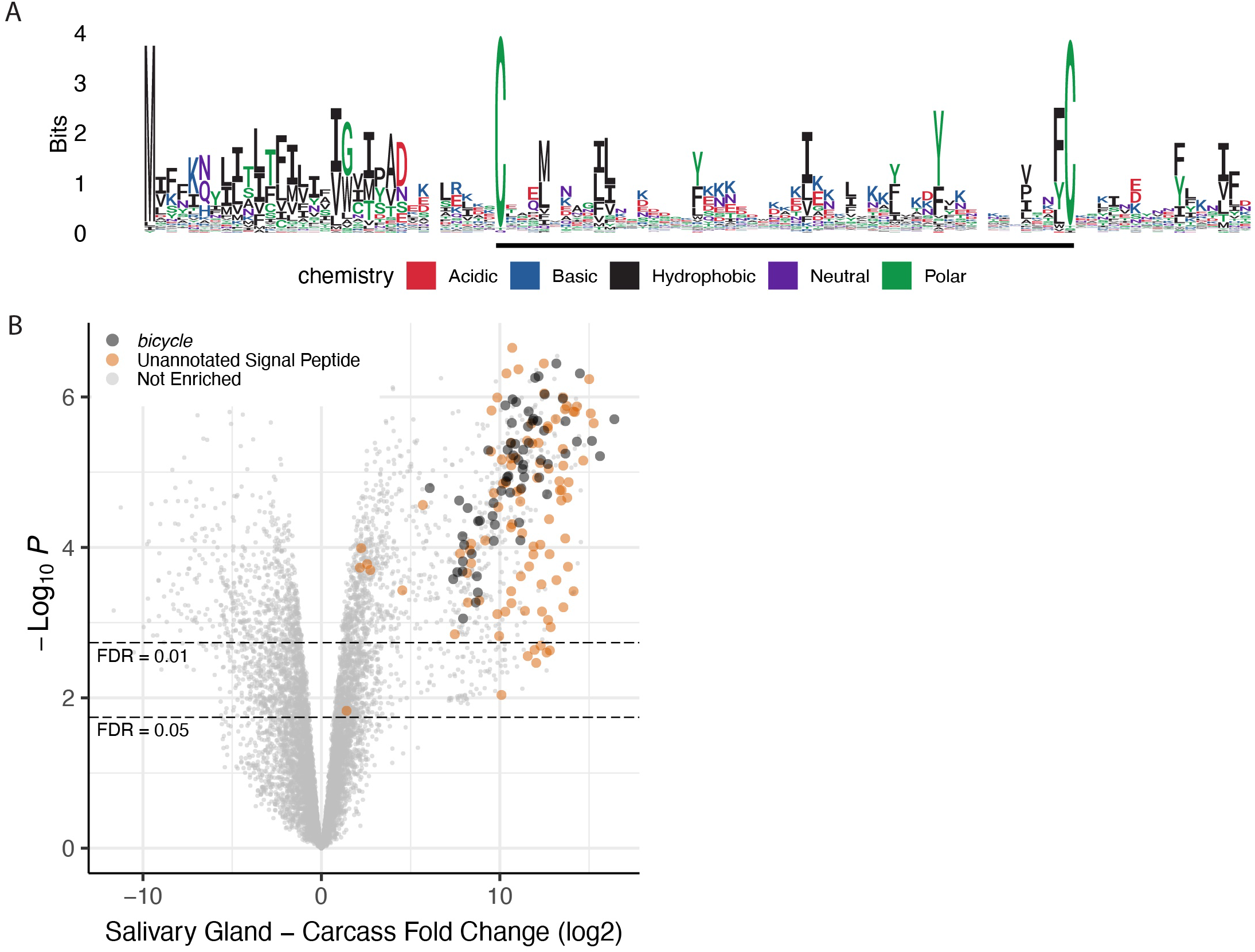
*A. pisum bicycle* homologs contain a single CYC motif and are strongly enriched in salivary glands. (A) Logo plot of *A. pisum bicycle* homologs detected by gene-structure based classifier. Most *A. pisum* homologs contain a single CYC motif. (B) Volcano plot of differential expression between salivary glands and carcass reveals that *A. pisum bicycle* homologs are strongly over-expressed in salivary glands.

To explore whether these *A. pisum bicycle* homologs may contribute to the effector gene repertoire of *A. pisum*, we performed RNA-seq of salivary glands and carcasses. We found that 55% of the putative *bicycle* genes are among the most strongly overexpressed genes in salivary glands, along with a similar number of additional unannotated genes with signal peptides (Figure 5B). In addition, 17 proteins encoded by genes with multiple microexons were identified by a previously-published proteomic study of proteins secreted by *A. pisum* into an artificial food medium (Dommel et al., 2020), and we found that at least 13 of these proteins are *bicycle* homologs (Table S1). In summary, the *A. pisum* genome encodes approximately 121 *bicycle* homologs that may contribute to the *A. pisum* effector protein repertoire.

### Bicycle genes are present in genomes of aphids, phylloxerids, and coccids

The presence of *bicycle* homologs in three divergent aphid species, *H. cornu*, *T. ulmi*, and *A. pisum* implies that *bicycle* genes were present in the common ancestor of aphids, which lived approximately 280 MYA. To further explore the origins and evolution of *bicycle* genes, we downloaded genomes and RNA-seq data for nine additional aphid species and ten outgroup species from NCBI. We annotated predicted genes in all genomes using the same bioinformatic pipeline that we have used previously to discover *bicycle* genes in other species, applied the *bicycle* gene classifier to all predicted genes, and manually annotated all putative *bicycle* homologs guided by RNA-seq data. One caveat of this analysis is that many of these genomes are fragmented into many contigs and genes bridging contigs are often misannotated or not annotated, resulting in possible under-counting of *bicycle* homologs. In addition, we detected multiple putative *bicycle* homologs near the ends of contigs, suggesting that parts of single *bicycle* homologs are present on multiple contigs, which would lead to over-counting *bicycle* gene homologs. While these problems with current annotations are expected to increase variance in estimates of *bicycle* homologs, we did not detect a dependence of number of *bicycle* homologs detected on genome assembly quality (Figure S9A). In addition, we employed chromosome-level genome assemblies for two outgroup species where no predicted *bicycle* genes were found, *P. venusta* (Y. Li et al., 2020) and *T. vaporariorum* (Xie et al., 2020), suggesting that the failure to identify *bicycle* genes in these species did not result from poor genome assemblies.

We estimated the phylogeny for these 22 species using whole-genome proteomic predictions (Emms, 2019), and this phylogeny is in general agreement with earlier studies based on a small number of genes (Johnson et al., 2018; Nováková et al., 2013; von Dohlen and N. A. Moran, 1995) (Figure 1). We detected putative *bicycle* homologs in all aphid species studied here, supporting the inference that *bicycle* genes were present in the common ancestor of aphids (Figure 1). We found extensive variation in the number of *bicycle* genes between species. In all aphid species with a sufficient number of *bicycle* homologs, multiple sequence alignment revealed the presence of CYC domains (Figure S9B-E). In most Aphidini species (e.g. *M. persicae*, *A. pisum*, *P. nigronervosa*), most homologs appear to be *unicycle* genes (Figure S9B- C). In *Cinara cedri*, many homologs appear to be *tetracycles* (Figure S9E).

We found 69 putative *bicycle* homologs in *Daktulosphaira vitifoliae* indicating that *bicycle* genes were present in the common ancestor of the Aphidomorpha (Aphidoidea + Phylloxeridea). We also found 74, 26 and 2 putative *bicycle* homologs in the coccid species *Maconellicoccus hirsutus*, *Phenacoccus solenopsis*, and *Ericerus pela*, respectively. In *M. hirsutus* and *E. pela* profile *hmmer* search with the classifier identified genes identified 17 and 12 additional candidate *bicycle* genes, respectively. The putative *bicycle* homologs from these non-aphid species do not contain obvious CYC motifs (Figure S10A-C). However, gene structure aware alignment with the *H. cornu bicycle* genes revealed that the putative homologs from *D. vitifoliae*, *M. hirsutus*, and *P. solenopis* share more introns in the same locations with *H. cornu bicycle* genes (Figure S10D-F) than expected by chance (Figure S6E), supporting the inference that these are *bicycle* homologs.

We did not detect *bicycle* homologs in any of the species sampled from the Psylloidea, Aleyrodoidea, or Fulgoroidea. These results indicate that *bicycle* genes were present in the common ancestor of coccids and aphids and may have evolved in the common ancestor of these lineages. However, for at least two reasons we cannot rule out the possibility that *bicycle* genes originated earlier. First, we observed extensive variation in the number of *bicycle* homologs across lineages, so *bicycle* genes may have been lost in the specific outgroup species studied here. Second, ancestral *bicycle* genes may not display the specific gene structure detected by our classifier. Deeper taxonomic sampling and other homology search approaches may reveal an older origin of *bicycle* genes.

### Aphid genomes contain a conserved *megacycle* gene

In many of the aphid genomes we studied, our classifier identified a single extremely large gene containing more than 100 microexons and encoding a protein of approximately 2000 amino acids as a candidate *bicycle* homolog (Figure 6A). Since this gene was so large and most existing aphid genomes are incompletely assembled, we observed that many of these gene models were incomplete. We therefore performed *de novo* transcript assembly for all species studied using TRINITY (Grabherr et al., 2011) and manually assembled consensus transcripts of these long candidate *bicycle* genes.

**Figure 6.**
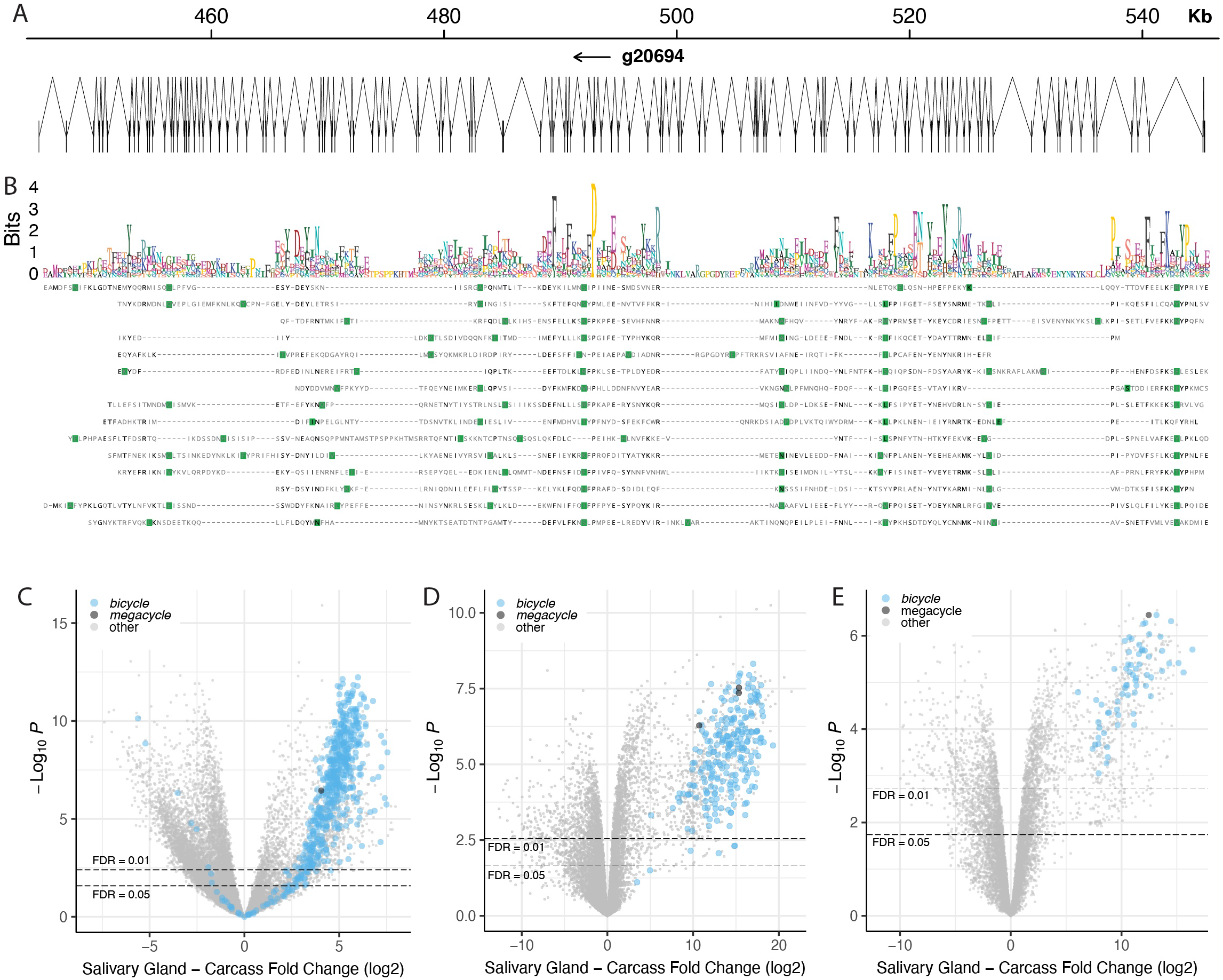
*Megacycle* genes have a repeating protein structure and are overexpressed in salivary glands of multiple aphid species. (A) Gene structure of the *M. sacchari megacycle* gene (*g20694*). The gene includes 118 exons encoded in approximately 100 kbp of DNA. The protein is 2092 AA long. (B) Rapid Automatic Detection and Alignment of Repeats (RADAR) analysis of the *M. sacchari megacycle* gene provides evidence for 15 repeating units within the protein encoded by *g20694*. Multiple sequence alignment of these repeat units (not using information on gene structure) reveals that many introns, marked in green, are located in identical or similar positions in the aligned sub-sequences. A logo plot of the aligned repeated sequences is shown above the alignment. There is no obvious evidence for CYC motifs, although the motif is approximately the same length as the proteins encoded by many *bicycle* genes. (C-E) Volcano plots of differential expression between salivary glands and carcasses for three species, *H. cornu* (C), *T. ulmi* (D), and *A. pisum* (E), reveals that *megacycle* genes (black dots) are strongly over-expressed in salivary glands of all three species. *Bicycle* genes are marked as light blue dots.

Since we had previously found that *bicycle* gene length is correlated with the number of encoded CYC motifs (Figure 3), we explored whether these genes encoded proteins with multiple repeating units. Using Rapid Automatic Detection and Alignment of Repeats (Heger and Holm, 2000) via EMBL-EBI (Madeira et al., 2019), we found that these large proteins contain an N-terminal secretion signal and many repeating motifs and each motif is approximately the size of two CYC motifs. This structure was most obvious for the *Melanaphis sacchari* gene (Figure 6A). Sequence-based alignment of 15 predicted repeats of the *M. sacchari* protein, which did not use information of intron positions, revealed multiple shared intron locations across repeats (Figure 6B). Thus, this protein appears to have evolved through multiple internal tandem duplications.

This large protein exhibited little obvious sequence similarity to *bicycle* genes, but gene-structure aware alignment revealed multiple apparently conserved intron positions shared between the internal duplications of the large gene from *M. sacchari* and *H. cornu bicycle* genes (Figure S11A), suggesting that they share a common ancestor.

These large genes appear to be divergent Bicycle proteins that have duplicated the core motif many times. We therefore call these *megacycle* genes. Compared to most *bicycle* genes, *megacycle* orthologs are relatively well conserved across aphids and a phylogenetic tree reconstructed from the inferred Megacycle proteins is similar to the aphid phylogeny estimated from the whole proteome (Figure S11B). Most aphid genomes we examined contain a single copy of this gene; we did not detect this gene in *Diuraphis noxia*, and we found three paralogs in both *Cinara cedri* and *Tetraneura ulmi*. No *megacycle* orthologs were found in any of the non-aphid species.

For the three species for which we have good salivary gland transcriptomic data, *H. cornu*, *T. ulmi,* and *A. pisum* the *megacycle* genes are strongly expressed in salivary glands (Figure 6C-E). Thus, *megacycle* genes may contribute to the effector protein repertoire in aphids. The relatively conserved lengths and sequences of *megacycle* homologs suggests that they may share a relatively conserved role across aphids.

## Discussion

Some gene features, such as the structure of the encoded protein or the intron-exon structure of the gene, can provide evidence for homology that is independent of sequence (Bazan, 1991; Betts, 2001; Brown et al., 1995). This source of information is becoming increasingly available with the recent availability of many well-assembled, annotated genomes. It is possible to conceptualize a model that jointly utilizes both sequence and exon-intron structure in homology search, but we realized that *bicycle* genes are encoded with extremely unusual intron-exon structures and that a model incorporating only general features of exons may allow detection of distant homologs.

We found that a linear classifier using information on gene length, exon sizes, exon numbers, and exon phases provided a highly accurate predictor of *bicycle* homologs that could not be identified using sequence similarity searches. This predictor allowed identification of *bicycle* homologs across aphids and in several outgroup taxa (Figure 1). Homology was supported by *post hoc* observation of CYC motifs and of an excess of shared introns across many homologs. Thus, sequence independent features provide substantial evidence of homology and integration of even a few sequence independent features into existing sequence-based search methods may substantially increase power to detect ancient homology relationships between genes. All gene structure features that we used contributed to classifier performance, but no single feature was critically important for classifier performance. Therefore, in future work it would be ideal to incorporate multiple features of gene structure into homology search models.

For several reasons, our classifier may have underestimated the number of *bicycle* homologs in each species. First, this approach is sensitive to the quality of genome annotation, which is itself dependent on genome assembly quality. We found that most automatically annotated gene models of *bicycle* genes required manual re-annotation. Thus, we may have overlooked additional *bicycle* genes in these genomes because of genome fragmentation and inaccurate automated gene annotation.

Second, *bicycle* gene annotation in most species is likely hampered by the lack of salivary gland RNA-seq samples. We observed considerable variation in *bicycle* gene expression levels among samples, and deep sequencing of diverse samples is likely required to provide sufficient RNAseq evidence to build accurate *bicycle* gene models. Finally, it is possible that some *bicycle* homologs have evolved both divergent sequences and novel gene structures, preventing identification with any existing method.

### *bicycle* genes are not strictly associated with gall formation

*Bicycle* genes were detected originally as putatively secreted proteins strongly expressed in the salivary glands of the gall-forming aphid generation of *H. cornu* and variation in one *bicycle* gene is genetically associated with gall color (Korgaonkar et al., 2021). These observations led to the hypothesis that many *bicycle* genes participate in gall development. Our observations provide a revised assessment of potential Bicycle protein functions.

First, we found that many *bicycle* genes are most strongly expressed in salivary glands of generations of *H. cornu* that do not induce galls (Figure 2D-F). Second, two of the aphids studied here, *H. cornu* and *T. ulmi*, induce complex galls and their genomes include many more *bicycle* genes than other species we examined, but the genomes of two other gall-forming species, *E. lanigerum* and *D. vitifoliae* have only 27 and 69 *bicycle* genes, respectively. In addition, the genomes of some non-galling aphids have many *bicycle* genes, such as *Myzus persicae* and *A. pisum*, with 117 and 121, respectively. Thus, across species there does not appear to be a strong correlation between the number of *bicycle* genes and the gall-forming habit.

Given these new observations, we hypothesize that if Bicycle proteins have conserved molecular functions, then they likely have different targets in different plant species. In some cases, the targets may induce patterned cell proliferation, resulting in galls, and in others the targets may alter plant physiology in more cryptic ways to confer benefits on aphids.

None of the three coccid species we studied here induce galls. However, many coccids induce depressions, swellings, or other changes in plant organs (Beardsley and Gonzalez, 1975), including the induction of elaborate galls (Cook and Gullan, 2004). Parr (Parr, 1940) reported a remarkable series of experiments that appear to demonstrate that extracts of salivary glands of a coccid are sufficient to induce pit galls on oak trees. It may be worth exploring the hypothesis that *bicycle* genes contribute to the effector gene repertoire of scale insects and especially of the gall-forming species.

### New experimental opportunities provided by the presence of *bicycle* genes in many species

One of the challenges with studying the function of *bicycle* genes is that initially they were thought to be restricted to the non-model gall-forming insect *H. cornu*. *H. cornu* displays a complex life cycle, exhibiting many polyphenic forms and migration between two trees every year (Korgaonkar et al., 2021), and prospects for laboratory rearing and subsequent experimental manipulations of this species were poor.

The discovery of *bicycle* genes in other insects opens up the possibility that the function of *bicycle* genes can be studied in species more amenable to laboratory manipulation. For example, RNAi allows gene knockdown in *M. persicae* (Chen et al., 2020; Coleman et al., 2015; Gao et al., 2021; Mathers et al., 2017; Niu et al., 2019; Pitino et al., 2011) and *A. pisum* (Han et al., 2006; Mutti et al., 2008, 2006; Niu et al., 2019; Shakesby et al., 2009; Vellichirammal et al., 2017; Will and Vilcinskas, 2015) and CRISPR-Cas9 mutagenesis allows gene knockout in *A. pisum* (Le Trionnaire et al., 2019). Thus, the discovery of many bicycle genes in these species presents new experimental opportunities to uncover the organismal and molecular functions of *bicycle* genes.

## Methods

### Gene structure based bicycle gene classifier

The *bicycle* gene classifier was built as a generalized linear model (GLM) with the following predictor variables: total gene length, total length of coding exons, first coding exon length, last coding exon length, mean internal exon length, and number of internal exons in phase 0, 1, and 2. All length measurements are in base pairs. All genes with zero or one internal exon were removed from the training and test datasets because mean internal exon lengths and number of internal exons starting in each of the three phases cannot be accurately estimated, hence our classifier was unable to detect any potential *bicycle* genes with only one to three exons. The dependent variable from the GLM is the predictor of *bicycle* gene classification. The GLM was implemented with the *glm* function as *binomial* with *logit* link, and response variables for the whole-genome gene set were predicted using *predict*, both in the *R* base package *stats* v.3.6.1.

In the training set, the dependent variable of the GLM is whether the transcript was previously classified as a *bicycle* gene (Korgaonkar et al., 2021), where *bicycle* genes were coded as 1 and non-bicycle genes were coded as 0. We used only genes classified as either *bicycle* genes or previously annotated genes in the training set due to the possibility that unannotated genes include additional previously undetected *bicycle* genes. Previous annotation was defined as having at least one significant match at E < 0.01 after Bonferroni correction for multiple searches with *blastp* (*blast* v.2.9.0), *blastx* (*blast* v.2.9.0), or *hmmscan* (*HMMER* v3.2.1) against the Pfam database.

We evaluated the performance of the classifier using a precision-recall curve where 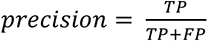 and 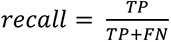, where TP, FP, and FN are the number of true positives, false positives, and false negatives, respectively. To determine a cutoff value on the dependent variable of the GLM for *bicycle* gene classification, we tested all possible cutoffs from 0 to 0.98 in increments of 0.02 to find the cutoff resulting in a precision to recall ratio closest to 1. We first validated our model by repeatedly sampling and training on 70% of the data and testing on the remaining 30% of the data for 100 replicates and found no substantial change to the trained classifier or the resulting cutoff value (mean 0.7338, s.d. 0.1578) amongst replicates.

We therefore trained the final classifier on all *H. cornu bicycle* genes and annotated non-*bicycle* genes without subsampling and determined a cutoff value of 0.72 (precision = 0.9925, recall = 0.9925). The coefficients to the independent variables in the GLM are as follows: intercept = 6.26, total exon length = -0.0206, total gene length = -0.0000642, first exon length = -0.00632, last exon length = 0.0103, mean exon length = -0.0655, number of exons in phase 0 = -3.32, phase 1 = -1.12, and phase 2 = 1.21. We evaluated the contribution of each independent variable by training the *H. cornu* dataset using GLMs with each term removed one at a time. We applied the GLM trained by the *H. cornu* dataset to twenty-one other species belonging to the families of Aphidoidae, Phylloxeroidae, Coccoidea, Psylloidea, Aleyrodoidea, and Fulgoroidea.

### Aphid collections and dissections

Aphids of *Tetraneura ulmi* were collected from galls collected on *Ulmus americana* on the grounds of Janelia Research Campus (Table S2-3). Aphids of *A. pisum* LSR1 were kindly provided by Greg Davis and grown on broad bean plants in the laboratory. Salivary glands were dissected out of multiple instars of fundatrices and their offspring for *T. ulmi* and of virginoparae for *A. pisum* and stored in 3 uL of Smart-seq2 lysis buffer (0.2% Triton X-100, 0.1 U/mL RNasin Ribonuclease Inhibitor) at -80°C for later mRNA isolation. Carcasses of both species were ground in PicoPure extraction buffer and mRNA was prepared using the PicoPure mRNA extraction kit. RNAseq libraries were prepared as described previously (Korgaonkar et al., 2021).

### Genome sequencing, assembly, and annotation of Tetraneura ulmi

HMW gDNA was prepared from *T. ulmi* aphids isolated from a single gall following the Salting Out Method provided by 10X Genomics (https://support.10xgenomics.com/genome-exome/sample-prep/doc/demonstrated-protocol-salting-out-method-for-dna-extraction-from-cells). 10X linked-read libraries were prepared and Chromium sequencing was performed by HudsonAlpha Institute for Biotechnology. Genomes were assembled with *Supernova* v.2.1.0 commands *run* and *mkoutput* (Weisenfeld et al., 2017).

We used 133,280,000 reads for an estimated raw genome coverage of 56.18X. The genome size of the assembled scaffolds was 344.312 Mb and the scaffold N50 was 10 Mb. Using BUSCO completeness analysis with *gVolante* version 1.2.0 and BUSCO_v2/v3 (Nishimura et al., 2017), we found that the genome contains 1035 of 1066 (97.1%) completely detected core arthropod genes and 1043 of 1066 (97.8%) partially detected core genes. The genome was annotated for protein-coding genes using BRAKER (Hoff et al., 2019) with multiple RNA-seq libraries (Table S3). This genome has been deposited to GenBank under the accession JAIUCT000000000.

### Differential expression analysis

Candidate *bicycle* genes from *H. cornu*, *T. ulmi*, and *A. pisum* identified by the classifier were manually re-annotated in APOLLO (Dunn et al., 2019) guided by RNAseq data viewed in IGV (Thorvaldsdóttir et al., 2013). Samples used for analysis are described in Table S3-5.

Differential expression analyses were performed as described previously (Korgaonkar et al., 2021) and full analysis scripts are provided at https://doi/figshare_spaceholder.

### hmmer and BLAST searches

The phylogeny of all species studied here was estimated using *Orthofinder* v 2.3.1 (Emms and Kelly, 2018) on the complete gene sets predicted by BRAKER for all genomes using the following settings: *-M msa -S diamond -A mafft -T fasttree*. The phylogeny was plotted using *ggtree* v. 2.0.4 in *R*.

We searched for *bicycle* homologs in all aphid species and outgroups using Position-Specific Interactive Basic Local Alignment Search Tool (PSI-BLAST) (Altschul, 1997) and *hmmer* (Eddy, 2011). We ran *PSI-BLAST* v. 2.9.0+ using *-max_target_seqs 5* and all other parameters as default. We used a cutoff of E < 0.01, Bonferroni corrected for 476 genes used as query searches. We ran profile-based *hmmsearch* v.3.2.1 using all default parameters and cutoff of E < 0.01 without multiple testing correction.

### Selection tests

*DnDs* between *H. cornu* and *H. hamamelidis* for all transcripts in the transcriptome was calculated using the *codeml* function from the *PAML* package v4.9j (Yang, 2007). The genome-wide selection scan was performed using *SweeD-P* v.3.1 (Pavlidis et al., 2013). Details of both analyses can be found in the Methods section of Korgaonkar et al., 2021.

### Gene-structure aware sequence alignment

We performed gene structure aware alignment using *prrn5* v 5.2.0 (Gotoh, 2021) with default weighting parameters. We prepared gene-structure-informed extended fasta files that were suitable as input to *prrn5* using the *anno2gsiseq.pl* utility provided at https://github.com/ogotoh/prrn_aln/blob/master/perl/anno2gsiseq.pl. We summarized the results of these multiple sequence alignments by generating histograms with a bin size of 1 to illustrate the fraction of sequences in the alignment that had an intron at each position in the alignment.

To determine whether intron positions were shared between genes more often than expected by chance, we computed the concordance in intron positions resulting from *prrn5* alignment of groups of unrelated genes. We identified paralog groups for the whole *H. cornu* transcriptome using *Orthofinder* v 2.3.3 (Emms and Kelly, 2018) and identified three paralog groups containing at least 10 genes and having at least 10 exons for comparison with *bicycle* genes: *SLC33A1*, *abcG23*, and *nrf-6*. Intron concordance was measured as the correlation coefficient (R) between the number of genes from the test gene family and the *bicycle* gene family containing an intron at each location in the alignment (Figure S6).

### Re-annotation of previously sequenced genomes

The *P. venusta* genome is an approximately full-chromosome genome assembly that was kindly provided to us by Yiyuan Li and Nancy Moran (Y. Li et al., 2020) prior to publication.

All other genomes, except for *H. cornu* and *T. ulmi*, were downloaded from NCBI (Table S5). We downloaded RNAseq data from NCBI (Table S4) and predicted coding genes using BRAKER (Hoff et al., 2019) with the same workflow we had used previously that had allowed discovery of *bicycle* genes (Korgaonkar et al., 2021).

## Availability of Data and Materials

All new sequence data generated during this study is available at NCBI at the accession numbers provided in Tables S2-4. All analysis scripts, which allow reproduction of all analyses and reproduction of all figures, are provided as Supplementary Material.

## Acknowledgements

We thank Andrew Lemire and the whole Quantitative Genomics team at Janelia Research Campus for generating and sequencing the RNA-seq samples, Tom Dolafi for maintaining our Apollo Annotation server, Zari Zavala-Ruiz and the Janelia Visitor Project for supporting CH, Fernando Bazan for recommending the use of RADAR software to detect internal repeats in *megacycle* genes, Gregory Davis for providing samples of *A. pisum*, and Patrick Reilly, Saskia Hogenhout and members of the Stern lab for critical feedback on this work and this manuscript, especially Aishwarya Korgaonkar.

## Contributions

DLS conceptualized the project, collected aphids, performed dissections, made DNA and most RNA samples, performed initial analyses, and wrote the paper. CH developed the bicycle gene classifier, independently verified all of the computational analyses, and assisted with preparations of figures and manuscript.

## Ethics declarations

HHMI has filed a provisional patent, number 63/243,904, for the inventors DLS and CH covering a gene structure, coding sequence-independent method of identifying bicycle genes.

## Supplementary Figures

**Figure S1.**
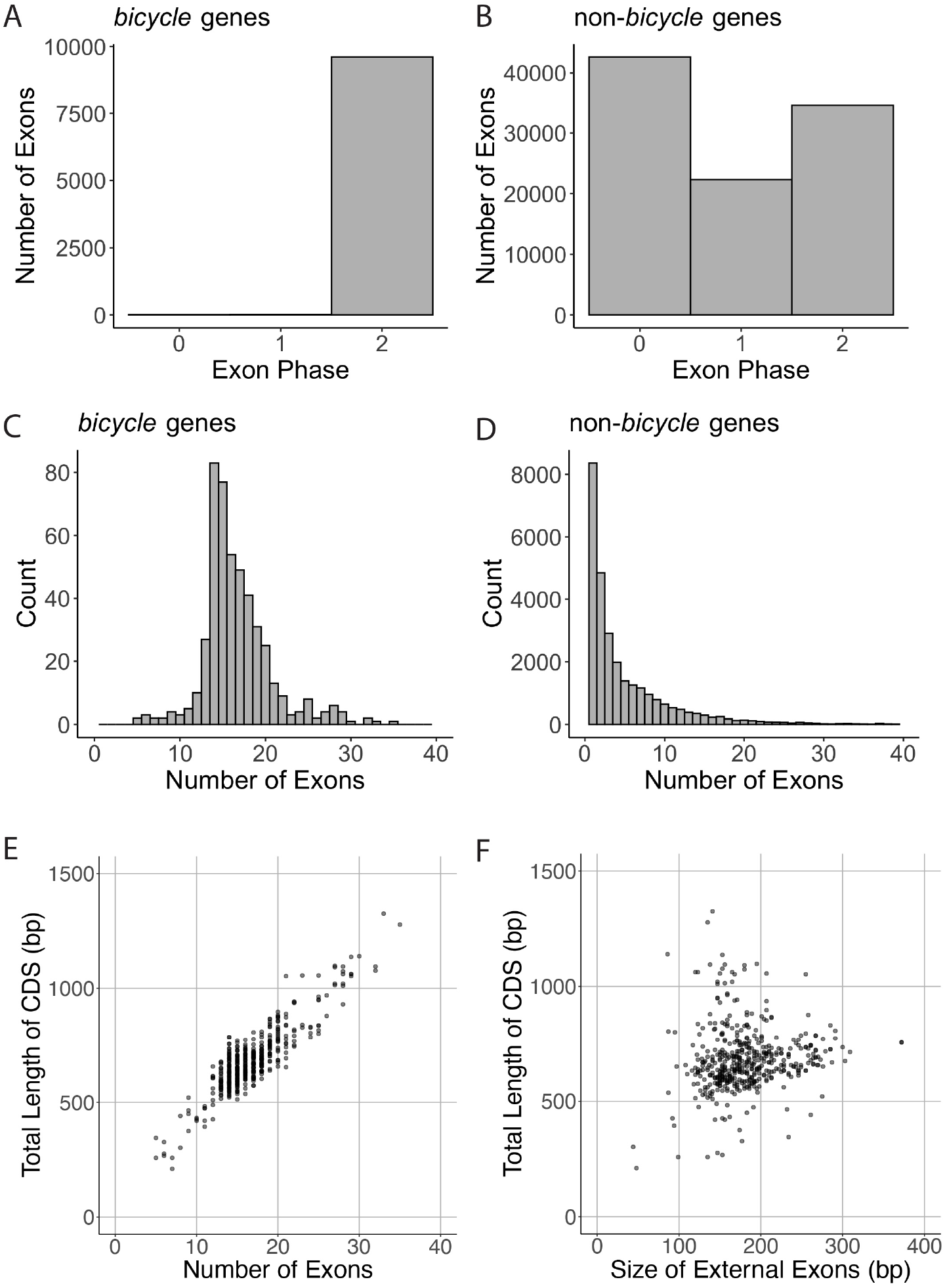
Salient features of gene structure of *H. cornu* bicycle genes. (A-B) Almost all internal exons of *bicycle* genes are of phase 2 (A), whereas the majority of internal exons of other genes in the *H. cornu* genome are of phase 0 (B). (C-D) *Bicycle* genes have a wide range of exon numbers, with a distribution (C) different from non-*bicycle* genes (D). (E-F) The total length of Bicycle proteins is better predicted by the number of exons in the gene (E) than by the length of the first and last exon (F).

**Figure S2.**
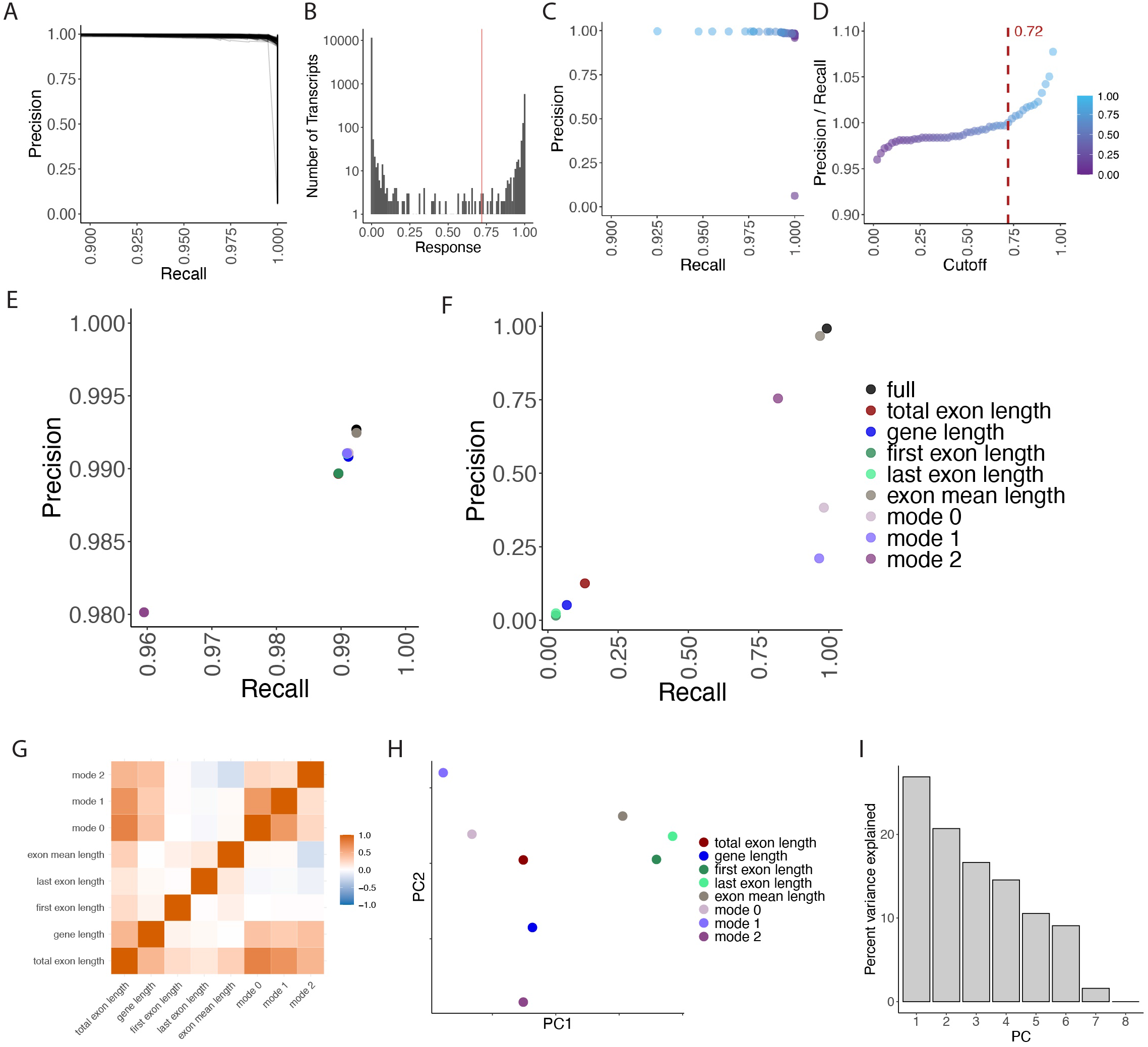
Performance of a gene-structure based classifier of *bicycle* genes. (A) Precision versus recall curves for replicate training runs of the classifier using 100 different subsets of the training data. The curves across all subsets are very similar, indicating that the classifier was not unduly influenced by a specific subset of the data. (B) Histogram of the number of transcripts with a particular classifier response shows a bimodal distribution. The vast majority of genes (>10,000) had a response close to 0. A threshold of 0.72 was chosen at a precision to recall ratio of 1. (C) Precision (the proportion of true positives among both true and false positives) versus recall (the proportion of true positives among true positives and false negatives) for the model trained on 476 previously identified *H. cornu bicycle* genes. Differently colored dots represent the precision to recall ratio for different probability cutoffs of the model, shown in panel (D). (D) The ratio of precision to recall for different cutoffs. Precision approximately equals recall (P/R = 1) at a cutoff of 0.72. (E) Precision and recall values for models where one variable at a time was removed from the model. Removal of the number of exons with mode2 had the largest decrement on performance, but all models performed well. (F) Precision and recall values for models where only one predictor variable was included in the model. Exon mean length had the best predictive value on its own, but no single variable performed as well as inclusion of all variables. (G) Correlation amongst predictor variables for all of the original 476 *bicycle* genes. Most variables are not strongly correlated. (H) First two principal components of principal component analysis of values for the predictor variables for all of the original 476 *bicycle* genes provides further support that variables are not strongly correlated. (I) Percent variance explained for the eight principal components of a principal component analysis for the predictor variables for all of the original 476 *bicycle* genes.

**Figure S3.**
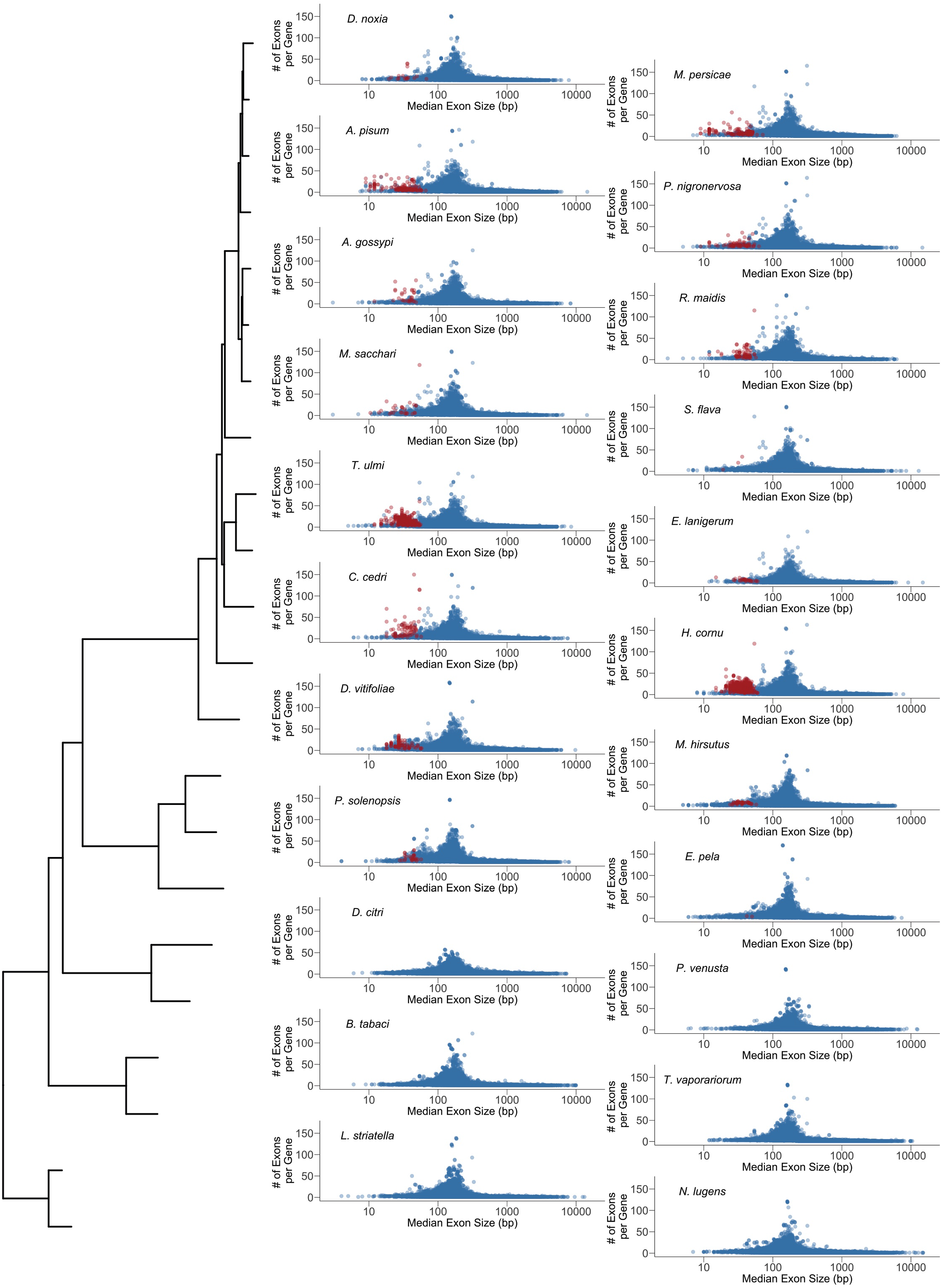
Plots of exon number per gene vs median exon size for all species studied. Phylogeny for all species is shown on left and is identical to phylogeny in Figure 1. *Bicycle* genes tend to have many small exons. Red = bicycle classifier, blue = other genes.

**Figure S4.**
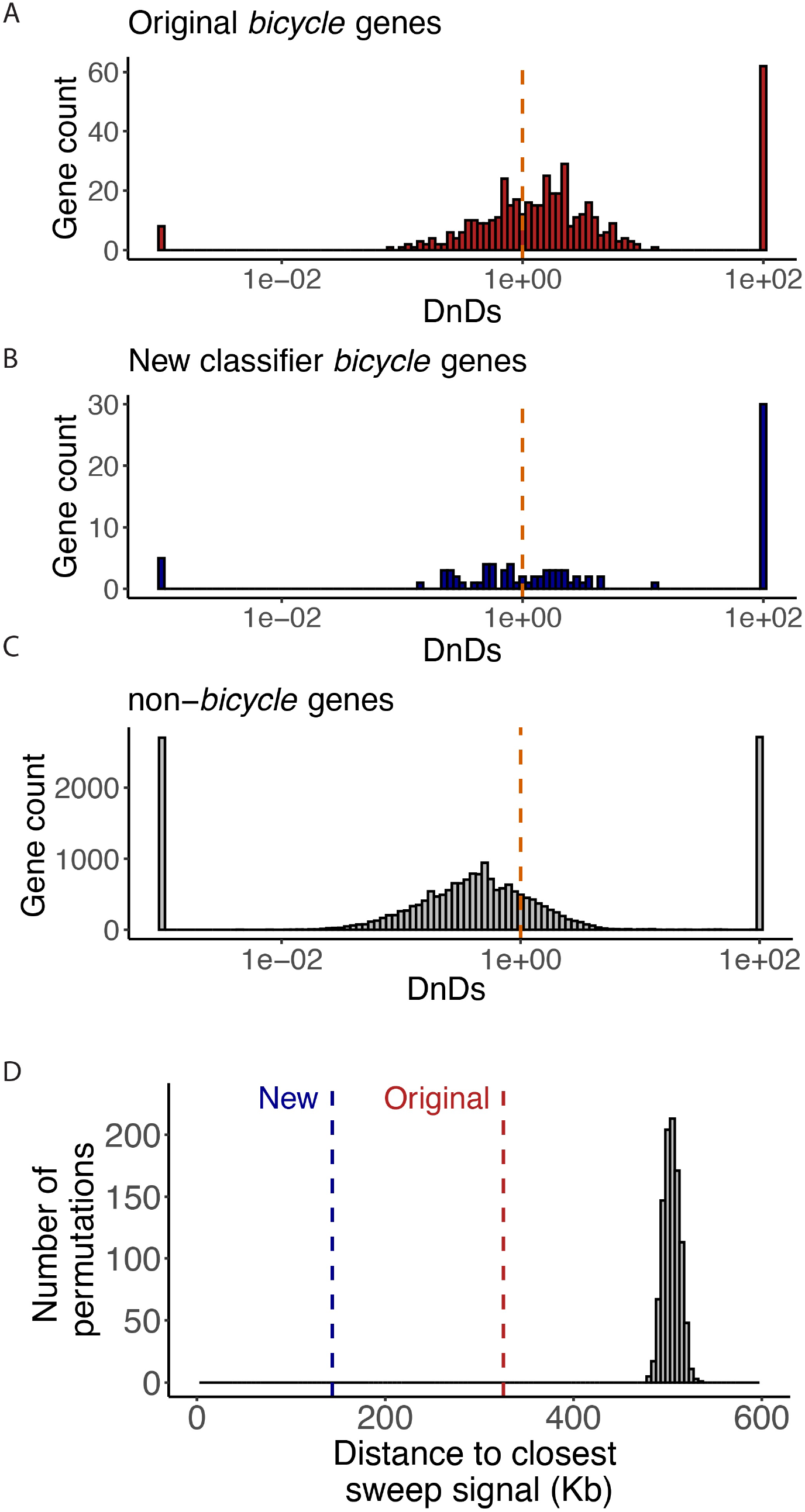
Newly identified candidate *bicycle* genes in *H. cornu* exhibit signatures of strong and recent positive selection. (A-C) *d_N_*/*d_S_* ratio for the 476 original *bicycle* genes (A), the newly identified putative *bicycle* genes (B) and the non-*bicycle* genomic background (C) between *H. cornu* and *H. hamamelidis*. Both the original *bicycle* genes and the newly putative *bicycle* genes show an excess of genes with a ratio of *d_N_*/*d_S_* above 1 compared to the background. Dashed vertical line indicates *d_N_*/*d_S_* = 1. (D) Median distance to the closest sweep signal for original *bicycle* genes (red dashed line) and classifier newly identified putative *bicycle* genes (blue dashed line). Histogram shows median distance to closest sweep signal for 1000 permutations of gene labels. The original *bicycle* genes and the new putative *bicycle* genes are both closer to sweep signals than expected by chance.

**Figure S5.**
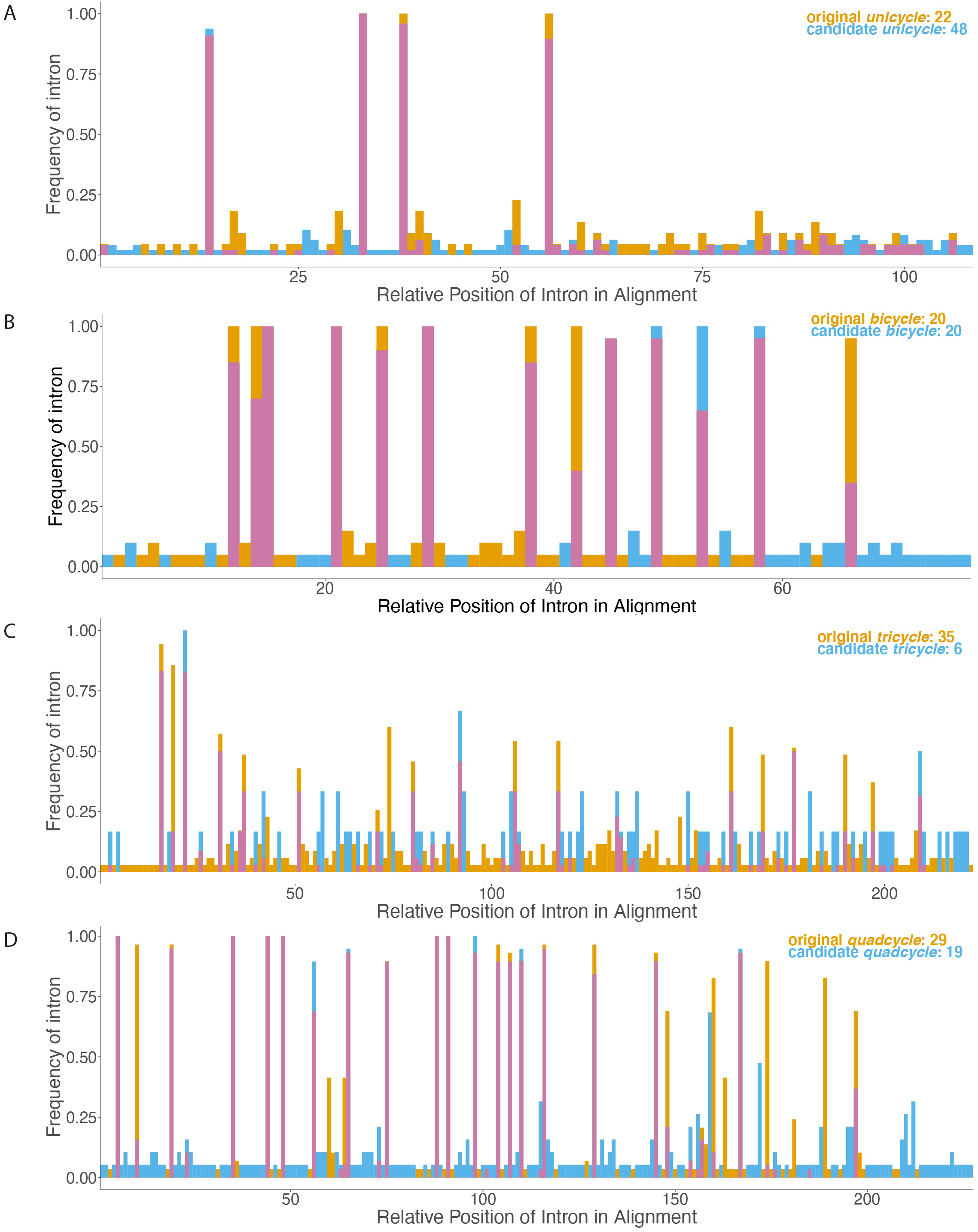
Gene-structure aware alignments reveal extensive conservation of intron locations between the originally-defined *bicycle* genes and putative *bicycle* homologs identified by the gene-structure based classifier. (A-D) Predicted proteins encoded by the original or newly identified bicycle homologs were divided by sequence length into Unicycle (A), Bicycle (B), Tricycle (C), or Tetracycle (D) proteins and aligned using gene-structure aware alignment. The frequency of introns at each location in the alignment is plotted for the original (orange) and newly defined (cyan) *bicycle* homologs and positions without introns were excluded from the plot. Intron positions are therefore relative to each other and do not indicate absolute position in the proteins or the alignments. Coincidence of intron positions is shown by purple bars. Original *bicycle* homologs were randomly down-sampled to 20 genes to match the number of newly identified *bicycle* homologs to improve overall alignment accuracy (B).

**Figure S6.**
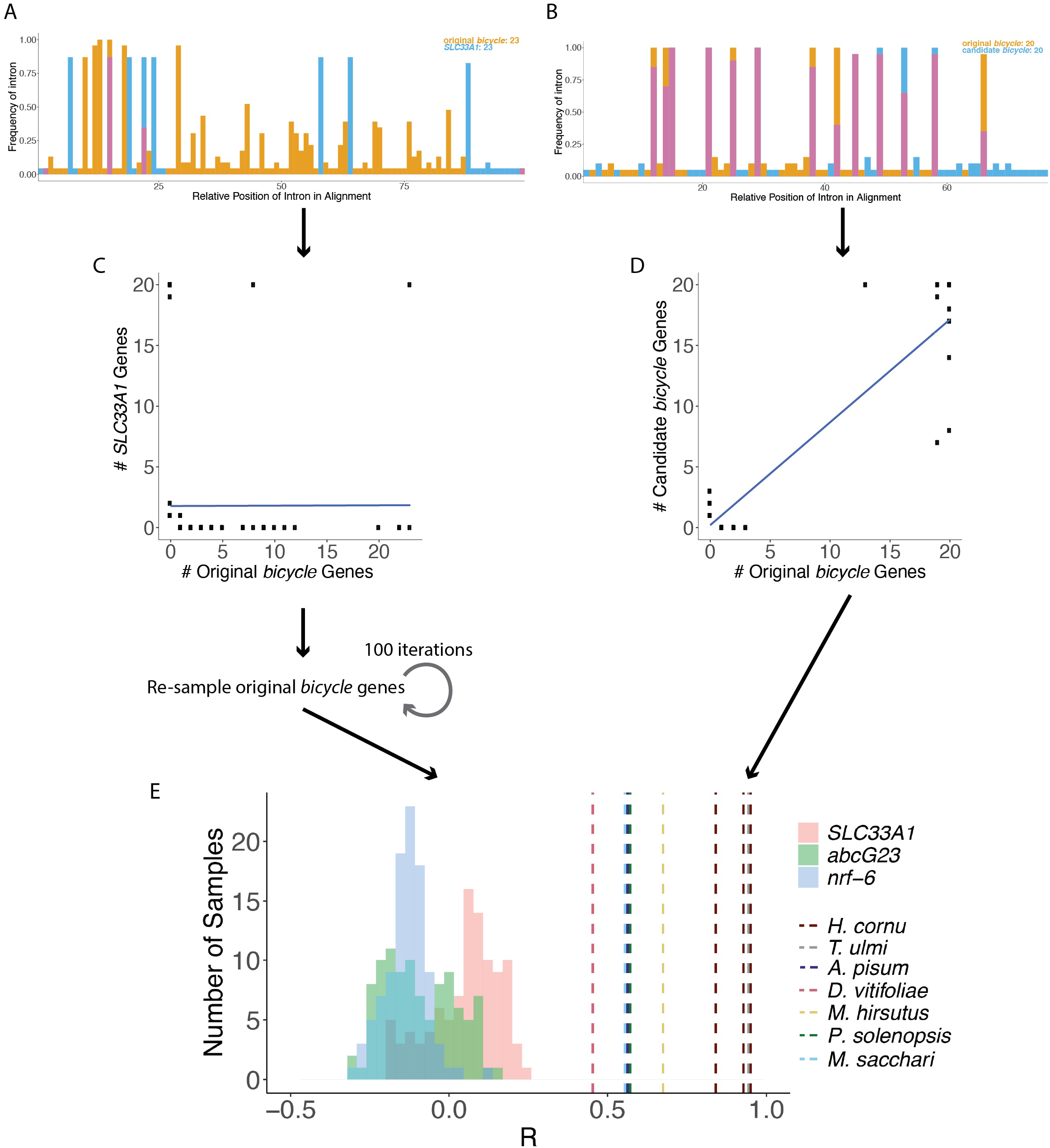
Testing intron concordance. To determine whether concordant introns were found more often than expected by chance alignment of unrelated genes, gene-structure aware alignment was performed between a subset of the original *bicycle* genes and genes from three gene families containing at least 10 genes with at least 10 exons each: *SLC33A1*, *abcG23*, and *nrf-6*. Intron concordance was measured as the correlation coefficient between the number of genes exhibiting an intron in the test gene family versus the original *bicycle* gene family. (A) Example of gene-structure aware alignment of 23 genes from *SLC33A1* (blue) and 23 original *bicycle* gene (orange) families. One intron was found frequently in both gene families (purple), a pattern observed in most alignments and likely reflects an artifact of the gene-structure aware alignment algorithm. (B) Example of gene-structure aware alignment of 20 candidate *H. cornu bicycle* genes identified by the classifier (blue) and 20 original *bicycle* genes (orange). Many introns are found in both groups (purple). (C-D) Examples of quantification of intron concordance, measured as the correlation between the number of genes with an intron at each location in the alignment in the test gene family versus the original *bicycle* genes. For a comparison of the unrelated gene families *SLC33A1* and original *bicycle* (C), the correlation was close to 0. For a comparison of candidate *bicycle* genes versus the original *bicycle* genes, the correlation was high and positive (D). (E) Intron concordance, measured as the correlation coefficient (*R*) of the number of genes sharing intron locations in gene-structure aware alignments, of repeated re-samplings of unrelated genes families generated a null distribution of expected *R* values for unrelated genes (*SLC33A1*, *abcG23*, and *nrf-6* versus original *bicycle* genes in red, green and blue histograms, respectively). *R* values for all comparisons of candidate *bicycle* genes found by the classifier and the original *bicycle* genes were all positive and much higher than any of the *R* values resulting from alignment of unrelated gene families, indicating that more introns are shared between genes in these gene-structure aware alignments than expected between unrelated gene families by chance (P < 0.01).

**Figure S7.**
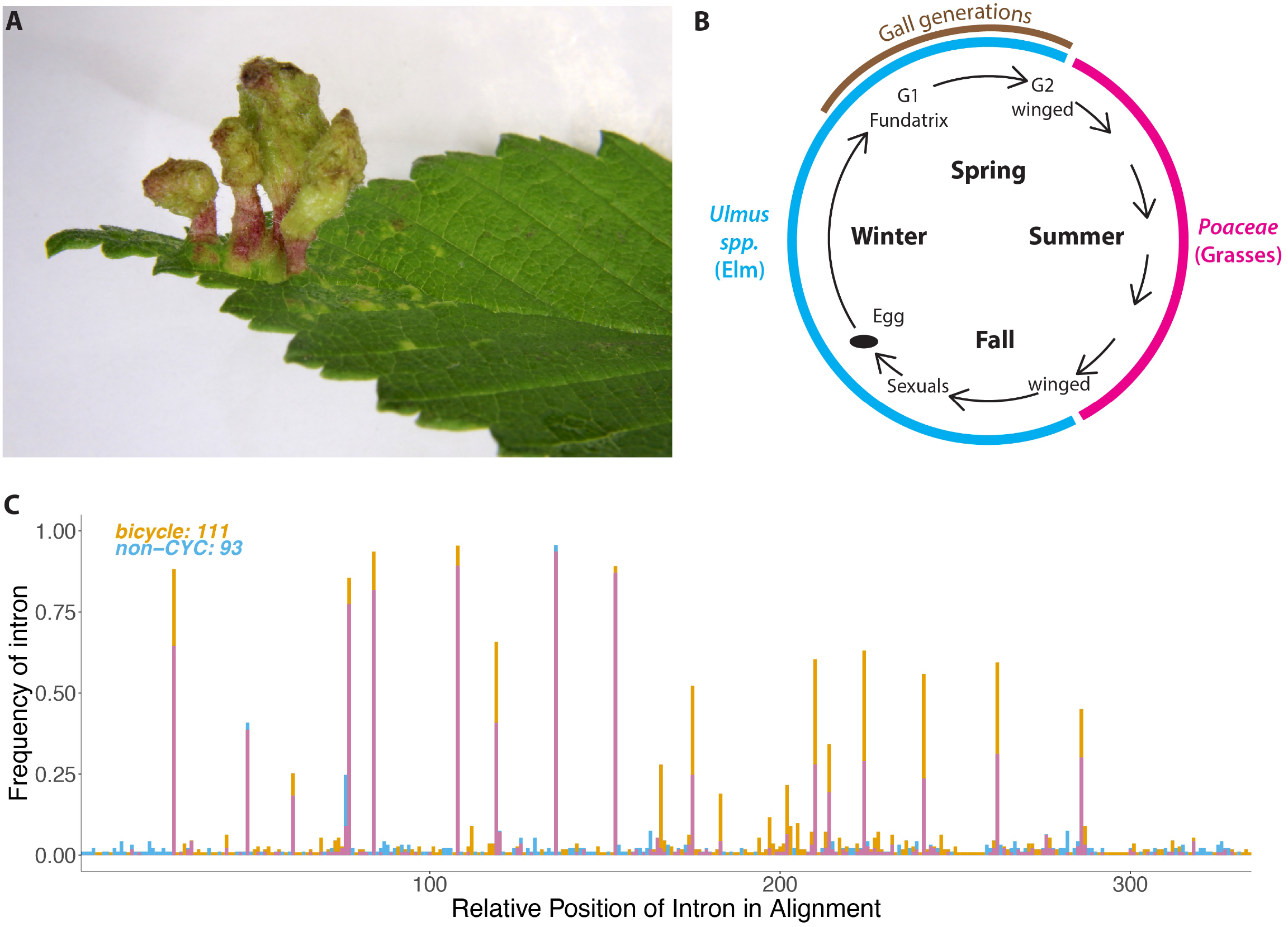
*Tetraneura ulmi* life cycle and additional features of *T. ulmi bicycle* genes. (A) Photo of mature galls of *T. ulmi* on leaf of *Ulmus americana* collected at Janelia Research Campus, Ashburn, VA, USA. (B) Diagram of life cycle of *T. ulmi*. Samples for differential expression analysis were collected from generations G1 and G2. (C) The *T. ulmi* putative bicycle homologs from Clusters 1 and 4, with and without CYC motifs, respectively, share multiple intron positions. Proteins from the two classes of genes were aligned using gene-structure aware alignment and the frequency of introns at each location in the alignment was plotted in different colors for the two classes of genes and positions without introns were excluded from the plot. Intron positions are therefore relative to each other and do not indicate absolute position in the proteins or the alignments. Sites of coincident introns are indicated in purple.

**Figure S8.**
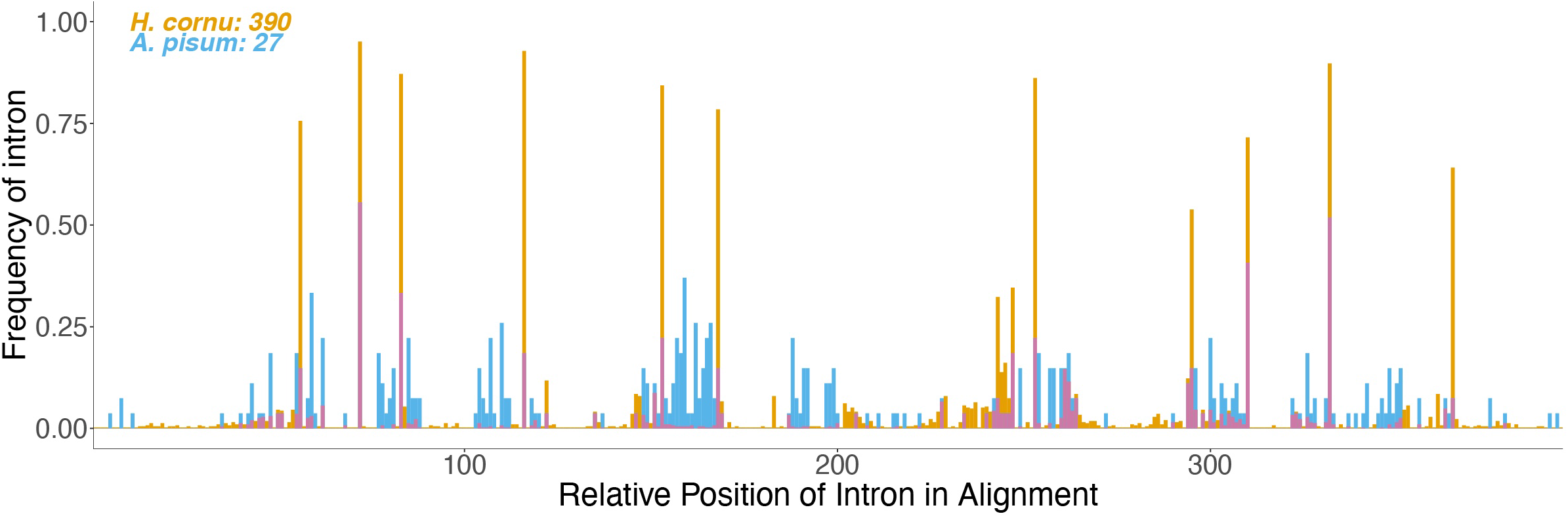
The *A. pisum* putative *bicycle* homologs share multiple intron positions with *H. cornu bicycle* genes. Proteins were aligned using gene-structure aware alignment and the frequency of introns at each location in the alignment was plotted in different colors for the two classes of genes and positions without introns were excluded from the plot. Intron positions are therefore relative to each other and do not indicate absolute position in the proteins or the alignments. Sites of coincident introns are indicated in purple.

**Figure S9.**
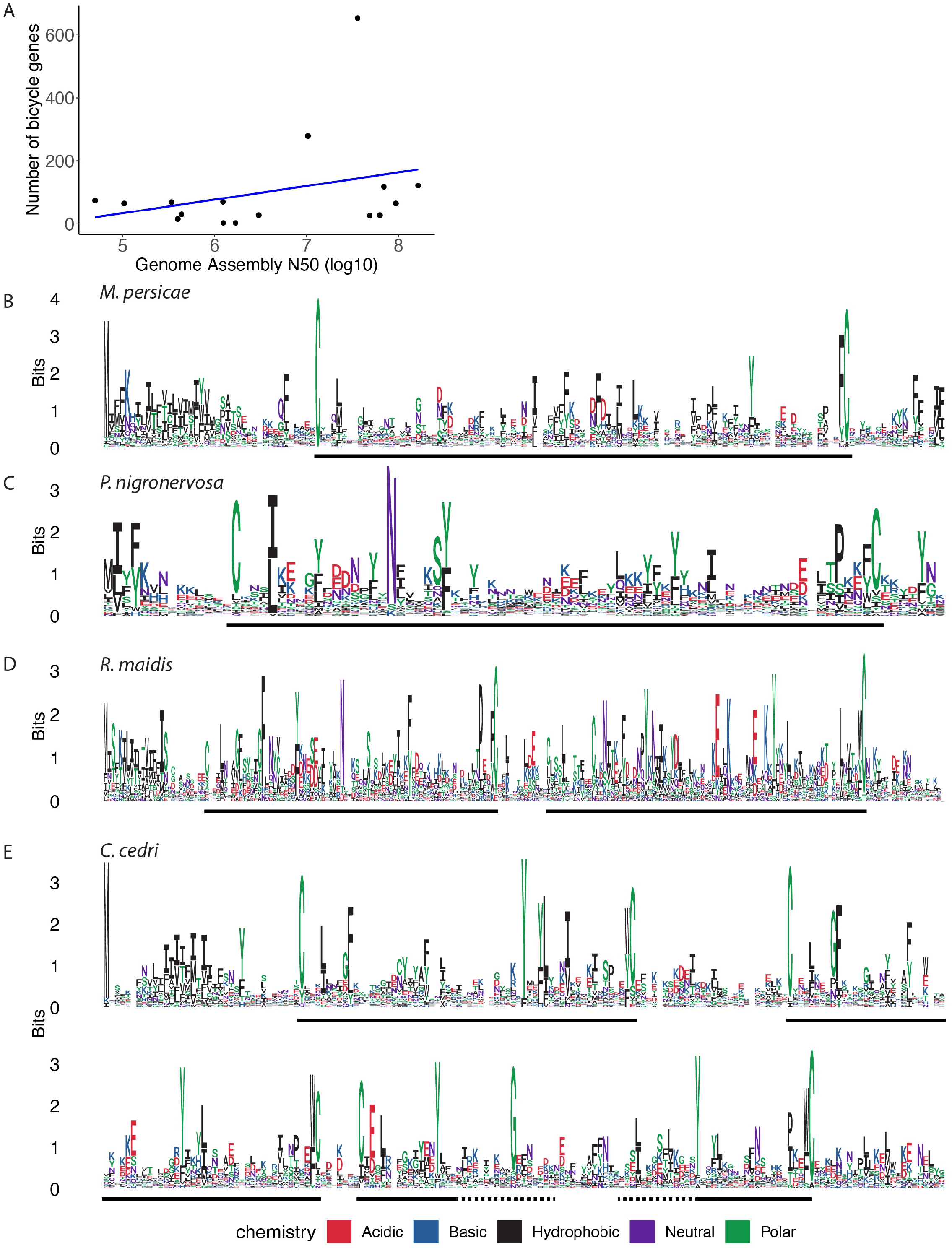
Features of *bicycle* homologs detected in aphid species. (A) A plot of the number of *bicycle* homologs detected by the gene-structure based classifier versus genome N50, a measure of genome assembly quality, shows no significant relationship between these variables (*R*^2^ = 0.09, *F* = 1.4, *P* = 0.26). (B-E) Logo plots of putative *bicycle* homologs detected in four aphid species, *M. persicae* (B), *P. nigronervosa* (C), *R. maidis* (D), and *C. cedri* (E) reveals that they all contain CYC motifs. The *M. persicae* and *P nigronervosa* genes are primarily *unicycles*, the *R. maidis* genes are primarily *bicycles*, and the *C. cedri* genes are primarily *tetracycles*.

**Figure S10.**
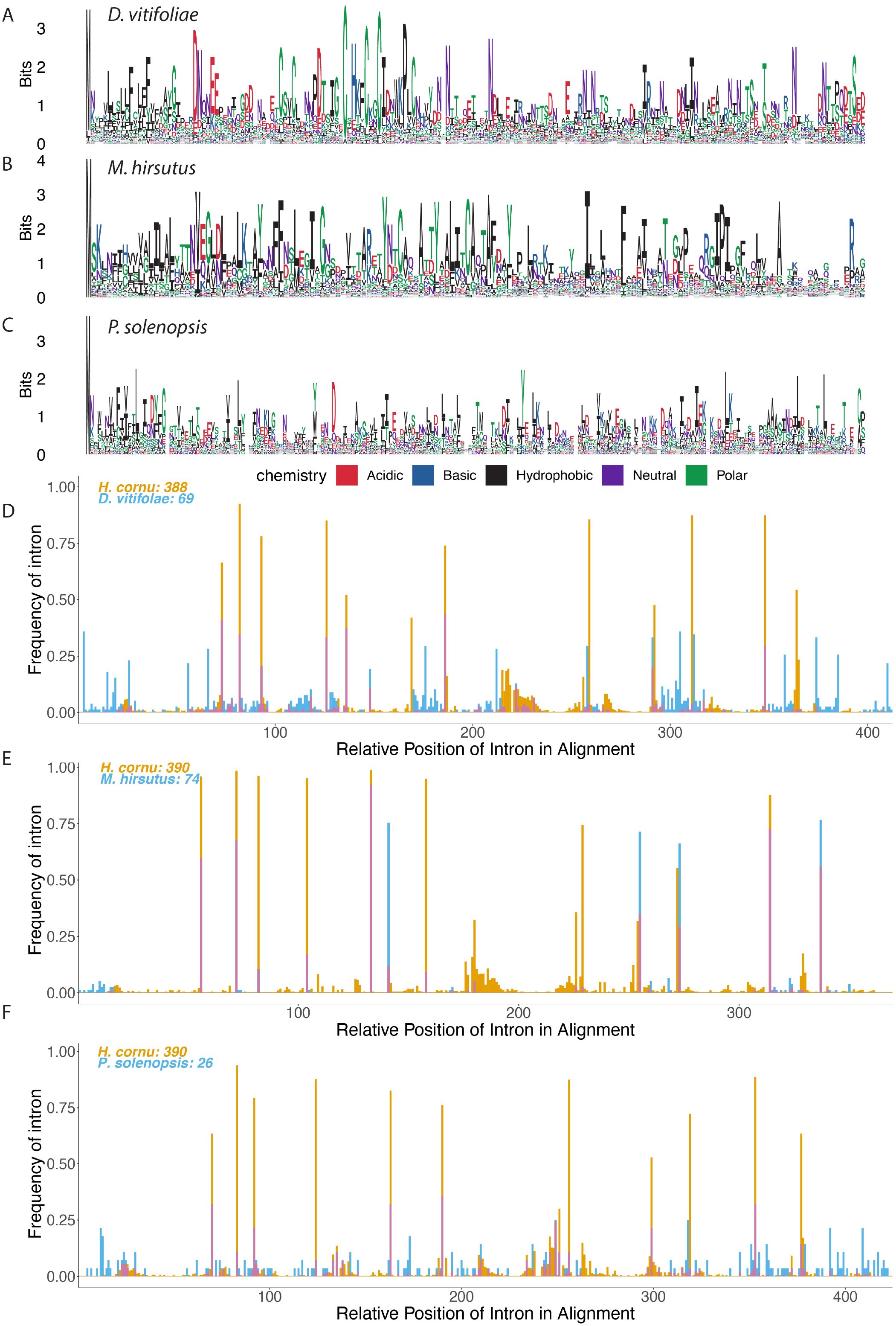
Features of *bicycle* homologs detected in outgroups to aphids. (A-C) Logo plots of proteins encoded by putative *bicycle* homologs detected in *D. vitilifoliae* (A), *M. hirsutus* (B), and *P. solenopsis* (C). The logo plots from these putative homologs show no obvious CYC motifs. (D-F) Gene-structure aware alignments reveal that multiple introns are shared between these highly divergent putative *bicycle* homologs and *H. cornu bicycle* genes. Positions in alignments without introns were excluded from the plot. Intron positions are therefore relative to each other and do not indicate absolute position in the proteins or the alignments. Sites of coincident introns are indicated in purple.

**Figure S11.**
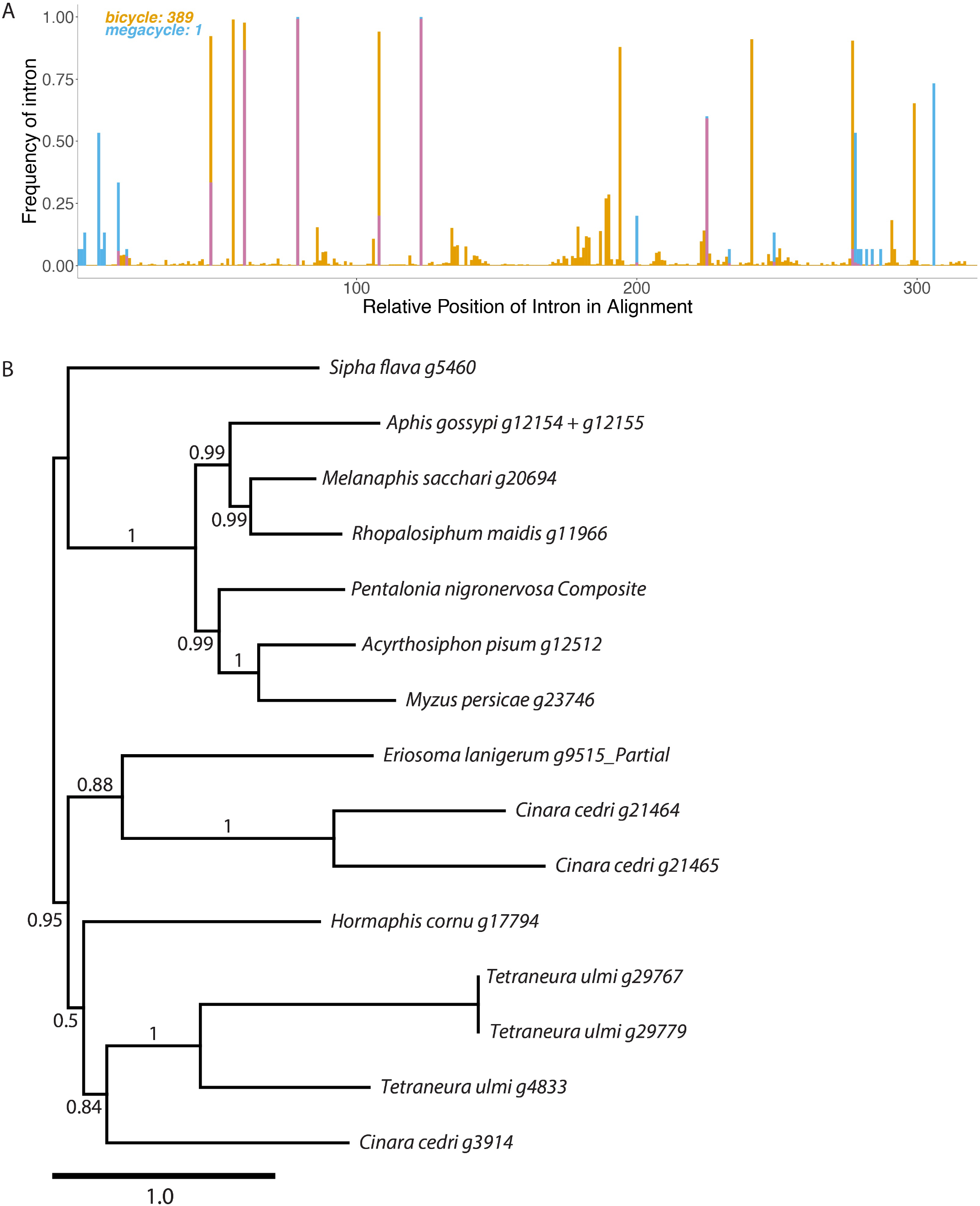
*Megacycle* genes share intron locations with *bicycle* genes and are found in many aphid species. (A) Gene-structure aware alignments reveal that multiple introns are shared between *megacycle* genes and *H. cornu bicycle* genes. Positions in alignments without introns were excluded from the plot. Intron positions are therefore relative to each other and do not indicate absolute position in the proteins or the alignments. Shared introns are shown in purple. (B) Maximum likelihood phylogeny of the protein sequences encoded by *megacycle* genes found in aphid species. *Megacycle* genes were not detected outside of aphids. The phylogeny has a topology similar to the aphid portion of the whole-proteome phylogeny shown in Figure 1. Only one *megacycle* gene was detected in most aphid species, but three paralogs were detected in *C. cedri* and *T. ulmi*. Values at nodes are bootstrap support values. Scale bar is 0.1 substitution per residue.

## Supplementary Tables

**Table S1.**
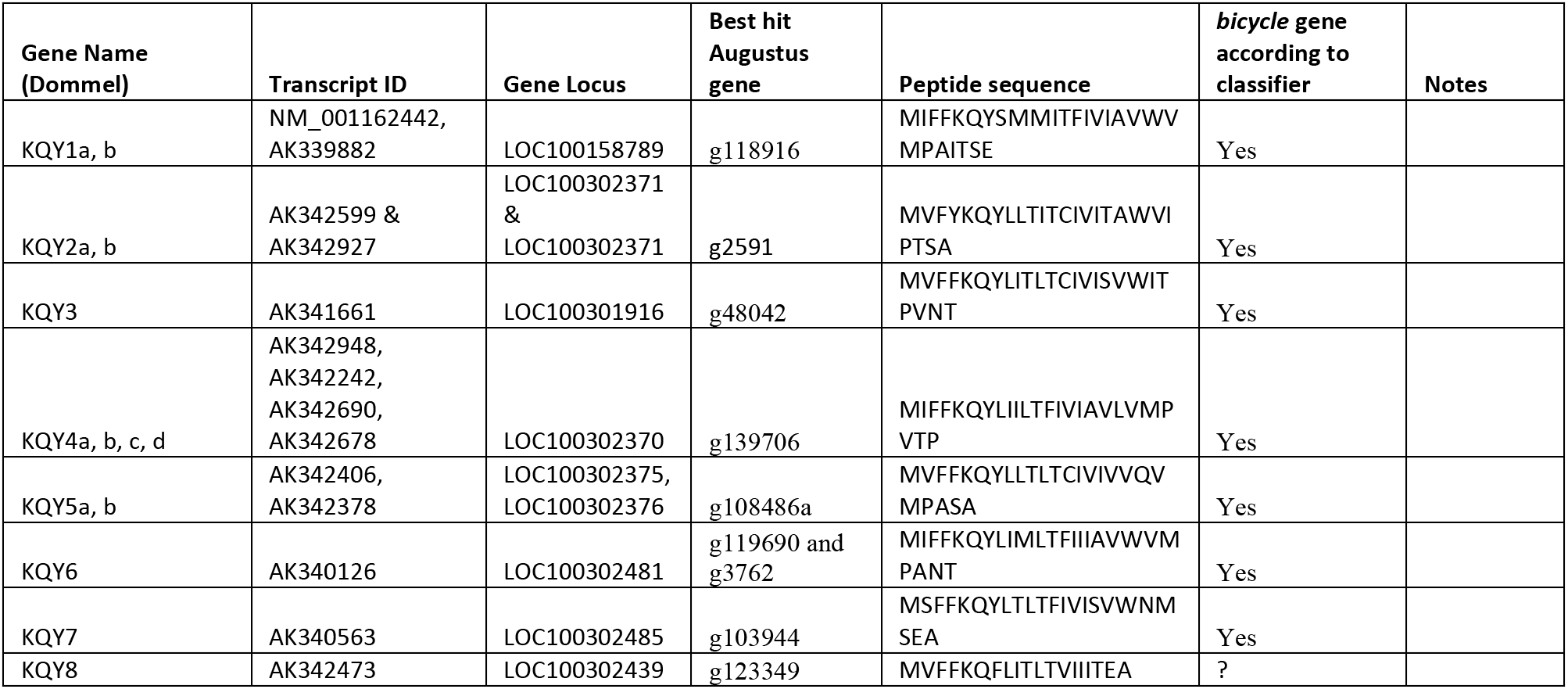
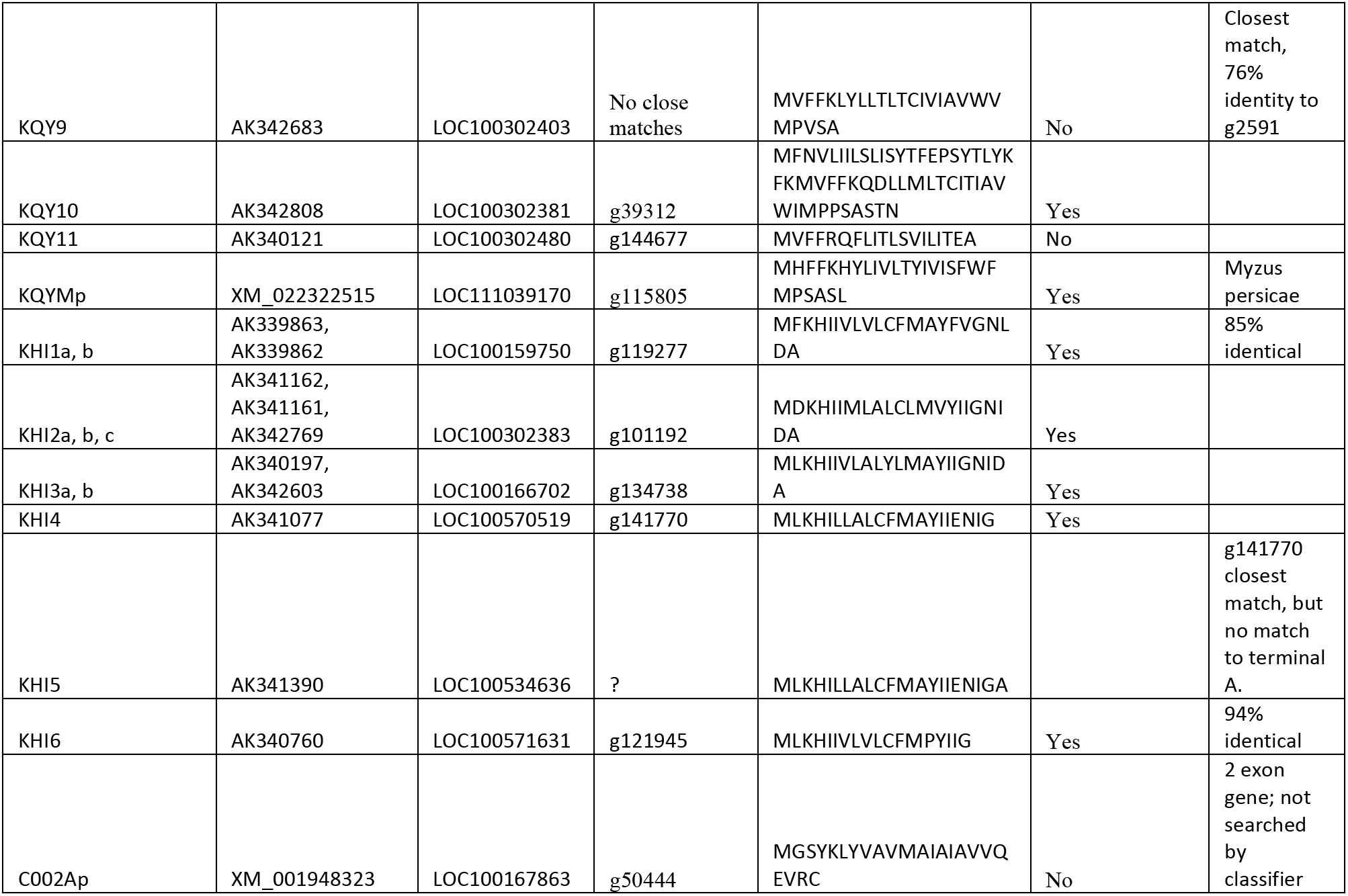
Many microexon genes producing secreted putative effector proteins in A. pisum detected by mass spectrometry of secreted proteins (Dommel et al., 2020) are identified as bicycle genes by the classifier.

**Table S2.**
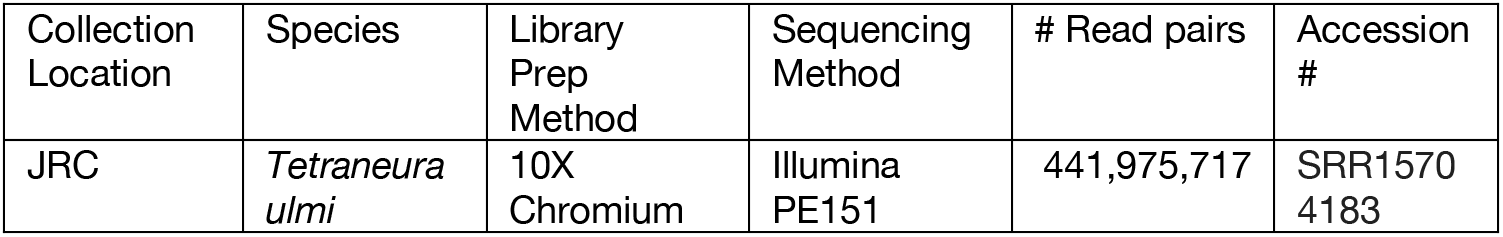
Details of Tetraneura ulmi sample, sequence files, and SRA accession numbers for genome sequencing. JRC refers to Janelia Research Campus, Ashburn VA, USA. These sequences can be found under the SRA BioProject PRJNA759586.

**Table S3.** Details of biological samples, sequence files, and SRA accession numbers for *Tetraneua ulmi* RNA-sequencing. JRC refers to Janelia Research Campus, Ashburn VA. These sequences can be found under the SRA BioProject PRJNA762862.

| Sample ID | Collection Date | Collection Location | Generation | Organ | Library Prep Method | Sequencing Method | # Reads | Accession # |
| --- | --- | --- | --- | --- | --- | --- | --- | --- |
| Tulm_Fund SG_2 | 4/22/19 | JRC | 1 | SG | WGB | Illumina PE150 | 8475508, 5308849 | SRR15871517 |
| Tulm_Fund SG_4 | 4/22/19 | JRC | 1 | SG | WGB | Illumina PE150 | 6087930, 6250323 | SRR15871516 |
| Tulm_Fund_SG_5 | 4/22/19 | JRC | 1 | SG | WGB | Illumina PE150 | 7474183, 7147731 | SRR15871 515 |
| Tulm_Fund_Carc_2 | 4/22/19 | JRC | 1 | Carcass | WGB | Illumina PE150 | 6208251, 6194859 | SRR15871 514 |
| Tulm_Fund_Carc_4 | 4/22/19 | JRC | 1 | Carcass | WGB | Illumina PE150 | 7453030, 6095776 | SRR15871 513 |
| Tulm_Fund_Carc_5 | 4/22/19 | JRC | 1 | Carcass | WGB | Illumina PE150 | 8399343, 7332616 | SRR15871 512 |
| Tulm_2ndGen_SG_2_1 | 4/22/19 | JRC | 2 | SG | WGB | Illumina PE150 | 6902051, 7037142 | SRR15871 511 |
| Tulm_2ndGen_SG_2_2 | 4/22/19 | JRC | 2 | SG | WGB | Illumina PE150 | 6888742, 6997123 | SRR15871 510 |
| Tulm_2ndGen_SG_2_3 | 4/22/19 | JRC | 2 | SG | WGB | Illumina PE150 | 6049169, 6696157 | SRR15871 509 |

**Table S4.** Details of biological samples, sequence files, and SRA accession numbers for Acyrthosiphon pisum RNA-sequencing. The aphid strain used was LSR1 (Caillaud et al., 2002), originally collected in the vicinity of Ithaca, NY, USA in summer 1998. Aphids were grown on broad bean plants in the laboratory at Janelia Reseach Campus and dissected for these samples on 21 February 2021. Sequences can be found under SRA BioProject PRJNA762703.

| Sample ID | Collection Date | Collection Location | Generation | Organ | Library Prep Method | Sequencing Method | # Reads | Accession # |
| --- | --- | --- | --- | --- | --- | --- | --- | --- |
| SG1 | 1998 | Ithaca, NY, USA | 1 | SG | WGB | Illumina PE150 | 1273082, 5724222 | SRR15862 168 |
| SG2 | 1998 | Ithaca, NY, USA | 1 | SG | WGB | Illumina PE150 | 1557673, 6606339 | SRR15862 167 |
| SG3 | 1998 | Ithaca, NY, USA | 1 | SG | WGB | Illumina PE150 | 1199959, 5221105 | SRR15862 166 |
| SG4 | 1998 | Ithaca, NY, USA | 1 | SG | WGB | Illumina PE150 | 1159521, 5169430 | SRR15862 165 |
| carc1 | 1998 | Ithaca, NY, USA | 1 | Carcass | WGB | Illumina PE150 | 998879, 4363153 | SRR15862 164 |
| carc2 | 1998 | Ithaca, NY, USA | 1 | Carcass | WGB | Illumina PE150 | 1185062, 5433749 | SRR15862 163 |
| carc3 | 1998 | Ithaca, NY, USA | 1 | Carcass | WGB | Illumina PE150 | 1045749, 4901408 | SRR15862 162 |

**Table S5.**
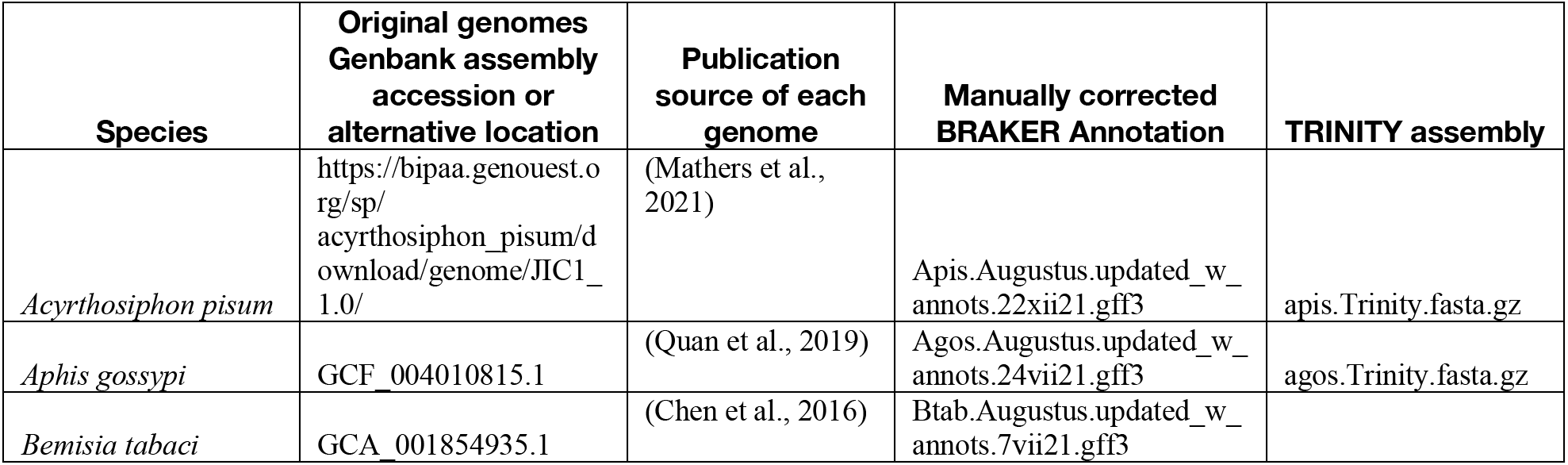
SRA accession numbers for Genomes downloaded from NCBI.

**Table S6.**
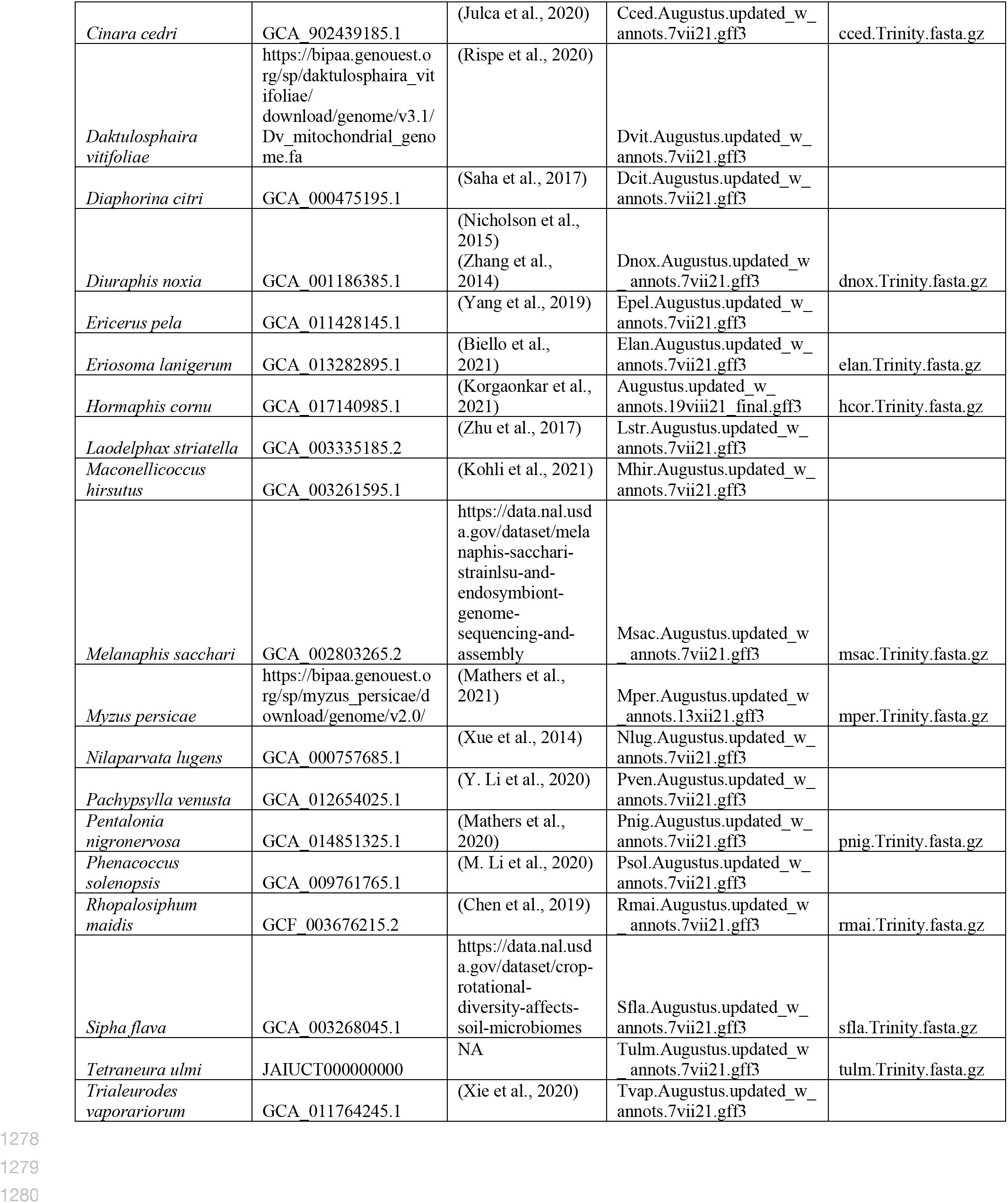
SRA accession numbers for RNAseq samples downloaded from NCBI. See separate file “S6_RNAseq_SRA_accessions.xlsx”.

